# CXXC5 is a ubiquitinated protein and is degraded by the ubiquitination-proteasome pathway

**DOI:** 10.1101/2024.11.06.622249

**Authors:** Hazal Ayten, Pelin Toker, Gizem Turan, Çağla Ece Olgun, Öykü Deniz Demiralay, Büşra Bınarcı, Gizem Güpür, Pelin Yaşar, Hesna Begüm Akman, Per Haberkant, Mesut Muyan

## Abstract

CXXC5 as a member of the zinc-finger CXXC family proteins interacts with unmodified CpG dinucleotides to modulate the expression of genes involved in cellular proliferation, differentiation, and death in physiology and pathophysiology. Various signaling pathways including mitogenic estrogens, particularly 17β-estradiol (E2), contribute to the expression and synthesis of CXXC5. However, how signaling pathways modulate protein levels of CXXC5 in cells is largely unknown. We previously reported that some key regulators, including retinoblastoma 1 and E74 Like ETS Transcription Factor 1, of the G1 to S phase transitions are involved in the expression of CXXC5 in estrogen-responsive MCF7 cells, derived from a breast adenocarcinoma. We, therefore, predict that the synthesis of CXXC5 is regulated in a cell cycle-dependent manner. We report here that although E2 in synchronized MCF7 cells augments both transcription and synthesis of CXXC5 in the G1 phase, CXXC5 protein levels are primarily mediated by ubiquitination independently of cell cycle phases. Utilizing the ^bio^Ubiquitination approach, which is based on cellular biotinylation of ubiquitin, in HEK293FT cells derived from immortalized human embryonic kidney cells followed by sequential immunoprecipitation coupled mass spectrometry analyses, we identified multiple ubiquitinated lysine residues of CXXC5. We show in both MCF7 and HEK293FT cells that the ubiquitinated lysine residues contribute to the degradation of CXXC5 through the ubiquitin-proteasome pathway.

## INTRODUCTION

17β-estradiol (E2), the main estrogen hormone in circulation, is a critical signaling molecule that contributes to the physiology and pathology of many organs and tissues, including breast tissue. The effect of E2 in epithelial cells of breast tissues is primarily mediated by the transcription factor, estrogen receptor (ER) α^1, 2^. Upon binding to E2, the activated ERα stably associates with estrogen receptor binding sites on DNA and regulates the expression of primary response genes, some of which encode enzymes for nucleic acid and protein metabolism, transcription factors, and membrane signaling proteins/receptors^3, 4, 5, 6^. Primary response gene products in turn participate in the expression of secondary response genes involved in the regulation of DNA synthesis, cell cycle, and cell division^3, 4, 5, 6^.

We^6^ and others^7^ reported that the CXXC5 is a primary estrogen-responsive gene. Also known as retinoid-inducible nuclear factor, RINF^8^, and WT1-induced Inhibitor of Dishevelled, WID ^9^, CXXC5 is a member of the ZF-CXXC family of proteins^10^. The CXXC5 gene is ubiquitously expressed with varying levels in human tissues^8,11^ http://atlasgeneticsoncology.org/gene/52549/). Evidence indicates that morphogenic retinoic acid^8^, multifunctional cytokine family member transforming growth factor-β^8^, bone morphogenetic protein BMP4^12, 13^, the Wnt family of secreted glycolipoprotein Wnt3a^14, 15, 16^ as well as Vitamin B2 and Vitamin D^17^ modulate *CXXC5* expression in a tissue-specific manner ^12, 16, 18, 19, 20, 21, 22^. Alterations in *CXXC5* expressions result in changes in cellular metabolism, proliferation, and/or differentiation in developmental processes and tissue maintenance^8, 12, 14, 18, 20, 21, 23, 24, 25, 26, 27, 28,29^. In keeping with the functional importance of CXXC5 in physiology, de-regulated expressions of *CXXC5* appear to contribute to various pathologies including Alzheimer’s disease, impaired wound healing in diabetes, acute myeloid leukemia (AML), gastric, prostate, and breast cancer^17, 30, 31, 32, 33, 34, 35, 36, 37, 38^.

Despite the importance of CXXC5 in diverse cellular events mediated by distinct signaling pathways in both physiology and pathophysiology, the regulation of *CXXC5* expression remains largely unknown. We recently explored the structural and functional features of the CXXC5 promoter as the key platform for the assembly of protein complexes to mediate transcription^39, 40^. We reported that *CXXC5* expression is driven by a core CpG island promoter which is associated with various transcription factors (TFs) and transcription co-regulatory proteins, as well as proteins involved in histone/chromatin, DNA, and RNA processing in cell models including E2-responsive and ERα-synthesizing MCF7 cells derived from a breast adenocarcinoma^41^. Of the possible TFs, we verified the association of the retinoblastoma protein (RB1) and E74-like ETS Transcription Factor 1 (ELF1) with the core *CXXC5* promoter and showed the involvement of ELF1 in the expression of *CXXC5*^41^. ELF-1 is a member of the ELF subfamily of ETS transcription factors that regulate many essential cellular processes, including cell cycle and proliferation^42^. ELF1 acts as an activator or repressor of target gene expressions by binding to a response element on DNA as well as through interactions with RB1^43^ which is a key regulator of the G1/S transition of the cell cycle^44^. Based on these together with our observations that CXXC5 contributes to the transition of E2 responsive and ERα synthesizing MCF7 cells synchronized at the G0/G1 phase to the S phase of the cell cycle critical for E2-mediated cellular proliferation^45^, we predict that *CXXC5* expression and/or synthesis is also regulated in a cell cycle-dependent manner.

We report here that although E2 augments both transcription and synthesis of CXXC5 in the G1 phase in synchronized MCF7 cells, CXXC5 protein levels are primarily regulated by ubiquitination independently of cell cycle phases. To identify the ubiquitinated lysine residues of CXXC5, we employed the ^bio^Ubiquitination (^bio^Ub) approach, which is based on the biotinylation of ubiquitin in cells^46, 47^. Our results from transiently transfected HEK293FT cells derived from immortalized human embryonic kidney cells using ^bio^Ub followed by sequential immunoprecipitation coupled-mass spectrometry analyses indicate that multiple lysine residues of CXXC5 are ubiquitinated. We find that the ubiquitinated lysine residues of CXXC5 contribute to the degradation of the protein through the ubiquitin-proteasome pathway in both MCF7 and HEK293FT cells.

## RESULTS

### Cell cycle-dependent expression and synthesis of CXXC5

The association of RB1 and ELF1 with the core promoter of *CXXC5* and the involvement of ELF1 in the regulation of *CXXC5* expression^41^ together with our observations that CXXC5 is a critical mediator of cell cycle regulation of MCF7 cells in response to E2-ERα signaling^45^ raise the possibility that CXXC5 expression and/or synthesis is regulated in a cell cycle-dependent manner. To address this issue, MCF7 cells were synchronized in G0/G1 by the use of the hormone-withdrawal approach we described previously^45, 48^. For this, we maintained MCF7 cells in the growth medium containing 10% charcoal dextran-treated FBS (CD-FBS) to remove steroid hormones including E2 for 72 h. Cells were then treated without (0.01% ethanol as the vehicle control) or with 10^-9^ M E2, an upper physiological level of circulating hormone in adult women^49^ for 2-6 h intervals up to 36 h. At each time point, collected cells were processed for flow cytometry (Fig 1A & 1B), RT-qPCR (Fig. 1C), and WB (Fig. 1D) analyses. Hormone withdrawal synchronized cells at the G0/G1 phase in which more than 80% of the cell population was accumulated. E2 treatment of cells effectively triggered cell cycle progression such that the proportion of cells began to transit into the S phase at 12 h reaching a plateau at 21 h (65 + 3.5%) in the S phase with corresponding decreases in the G0/G1 phase population (Fig. 1A & B). At subsequent time points, cells transited to the G2 phase, completing the cycle by 30 h resulting in a desynchronized cell population at 36 h. On the other hand, ethanol as control did not affect cycle phase distribution throughout the experimental periods; the majority (≥ 80%) of cells remained in the G0/G1 phase.

**Fig. 1.**
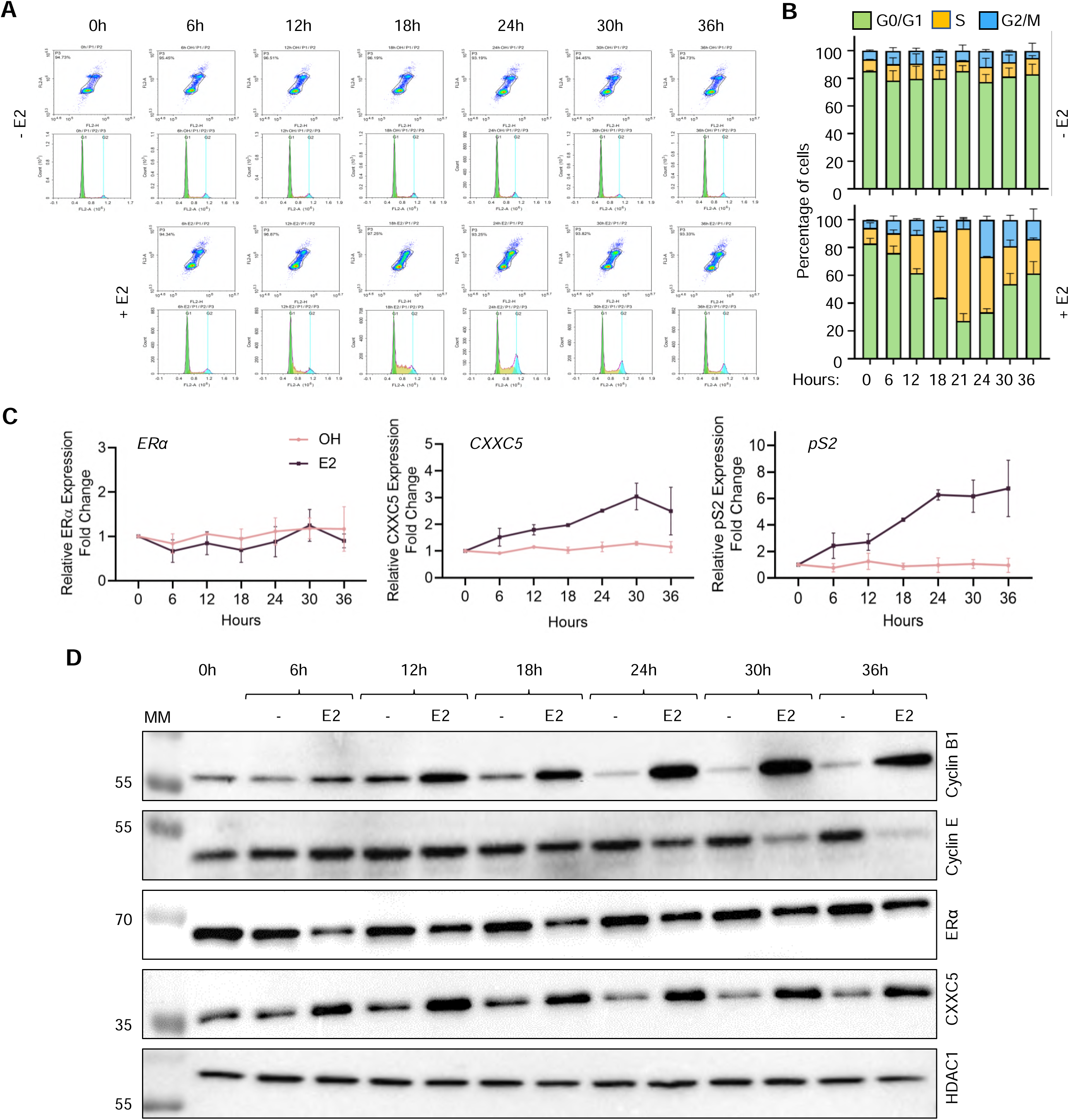
Cell cycle progression of MCF7 cells synchronized at G0/G1 by hormone withdrawal in response to E2. **(A-D)**MCF7 cells were grown in DMEM supplemented with 10% CD-FBS for 72 h with media change at 48 h. Cells were subsequently maintained in the same medium containing 0.01% ethanol (- E2) as vehicle control or 10^-9^ M E2 (+ E2) for 2-6 h intervals up to 36 h. At the termination, cells were collected with trypsinization. (**A & B**) A fraction of cells was subjected to the flow cytometry analysis and the remaining fractions were processed for (**C**) RT-qPCR and (**D**) WB analyses. Changes in the levels of CXXC5, TFF1, and ERα transcripts from three independent experiments were evaluated with RT-qPCR. Cellular extracts were also subjected to WB analyses for the modulation of protein levels of Cyclin B1 (Cyc B1), Cyclin E (Cyc E), ERα, CXXC5, or HDAC1 (as the loading control) using protein-specific antibodies. Representative images from the same experiment from flow cytometry and WB are shown. Molecular masses in kDa are indicated.

Transcript levels of ERα, which were normalized to the geometric means^50^ of the transcript levels of *RPLP0* (60 S acidic ribosomal protein P0) and *PUM1* (Pumilio RNA Binding Family Member 1), which are used as reference transcripts in breast carcinoma cell models^51^, were unaltered whether or not cells were treated with E2 (Fig. 1C). Whereas, E2 treatment of synchronized MCF7 cells steadily increased CXXC5 transcript levels reaching maximal levels by 24 h as similarly observed with the expression of *TFF1 (Trefoil Factor 1)*/*pS2*, a well-characterized E2-ERα responsive gene^45, 52, 53^ which we used here as a control (Fig. 1C).

Cell cycle progression triggered by E2 was reflected in changes in protein levels of Cyclin B1 (Fig. 1D), which is the regulatory subunit of cyclin-dependent kinase 1 (CDK1) essential for the transition from the G2 phase to mitosis^54^. With the E2 treatment, Cyclin B1 levels increased in the early S phase and remained elevated until the desynchronization of cells. On the other hand, Cyclin E as an activator of CDK2 (Cyclin-Dependent Kinase 2) critical for the entry to and progression through the S phase^55^ began to decrease at 24 h corresponding to the late S phase reaching low levels by 36 h (Fig. 1D). Diverging from ERα transcript levels, E2 treatment rapidly decreases the protein levels of ERα^56, 57^. Consistent with this, the E2 treatment of synchronized MCF7 cells for 6 h reduced ERα levels to a nadir which was maintained throughout the cell cycle until the desynchronization of cells at 36 h (Fig.1 D). In contrast, CXXC5 reached the highest levels by 4-6 h of E2 treatment and remained at similar levels in subsequent phases (Fig. 1D and Supplementary Information Fig. S1). HDAC1 levels used as the loading control were unaltered whether or not cells were treated with E2. A steady increase in CXXC5 transcript levels throughout cell cycle phases as opposed to the protein levels which rapidly increase and reach a plateau within 6 h post-E2 treatment corresponding to the early G1 phase implies post-transcriptional regulation of protein levels which could involve stabilities of transcripts and/or the CXXC5 protein.

### Examination of CXXC5 protein stability upon transcription and/or translation inhibition

We then addressed whether discordant changes in transcript and protein levels of CXXC5, as observed with ERα, throughout the cell cycle involve the stabilities of transcript and/or protein. To examine this issue, we examined the effects of blocking general transcription and/or translation on transcript and protein levels of CXXC5 with RT-qPCR and WB, respectively. To block transcription, we used actinomycin D (ActD) which intercalates into DNA thereby preventing the progression of RNA polymerases^58^. For the inhibition of protein synthesis, we used cycloheximide (CHX) which interferes with the translocation step in protein synthesis by blocking tRNA binding and release from ribosomes^59, 60^. To evaluate the minimal concentration as well as the pre-treatment duration of ActD and/or CHX in effectively blocking transcription and/or translation in MCF7 cells, we used 5-Ethynyl Uridine (5-EU) and/or L-homopropargylglycine (L-HPG) incorporation into nascent transcripts and/or polypeptide chains by chemoselective ligation followed by fluorescence microscopy (Supplementary Information, Fig. S2). Based on the results, we selected to use 5 µM ActD and/or 50 µg/ml CHX in our experiments. MCF7 cells synchronized in G0/G1 for 72 h by hormone withdrawal were treated without (0.01 % ethanol as vehicle control) or with 10^-9^ M E2 for 36 h to simulate a desynchronized cell population. We then incubated cells in a fresh growth medium without or with 10^-9^ M E2 in the absence or presence of 5 µM ActD and/or 50 µg/ml CHX for various durations (0, 0.5, 1, 2, 4, and 8 h). Collected cells were processed for and subjected to RT-qPCR and WB analyses (Fig. 2).

**Fig. 2.**
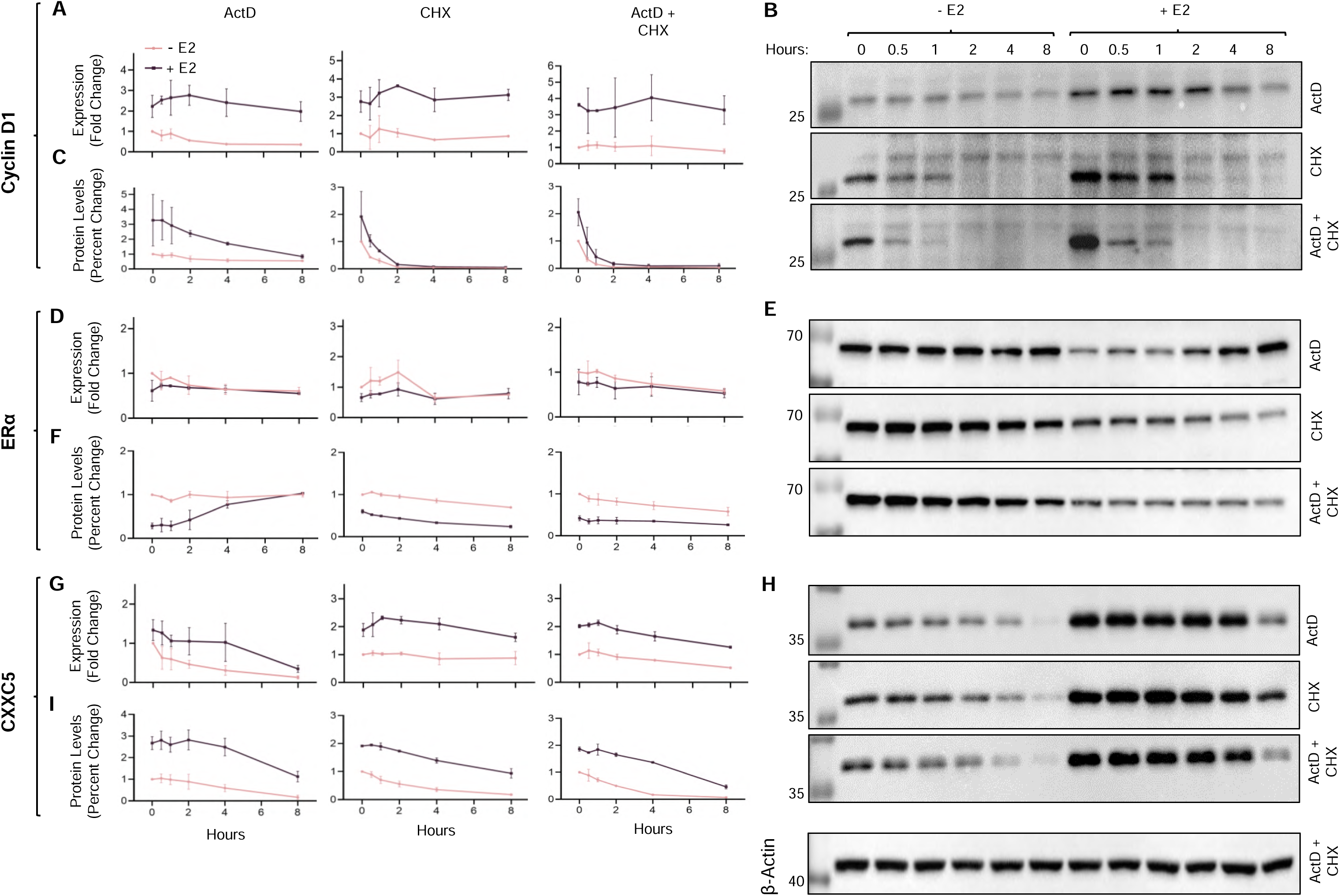
Stability of CXXC5 transcript and protein in the absence or presence of E2. Synchronized MCF7 cells for 72 h with hormone withdrawal were treated without (- E2) or with 10^-9^ M E2 (+ E2) for 36 h. Cells were then treated with 5 µM actinomycin D (ActD) for the assessment of transcript stability and/or 50 µM cycloheximide (CHX) for the examination of protein stability for 0, 0.5, 1, 2, 4, and 8 h in the absence (- E2; 0.01 % ethanol as vehicle control) or presence of 10^-9^ M E2 (+ E2). A fraction of cellular extracts was subjected to RT-qPCR using primers specific to (**A**) Cyclin D1, (**D**) ERα, or (**G**) CXXC5 mRNA. The remaining fraction was used for the visualization of WB and densitometric analysis relative to β-Actin levels as the loading control with the antibody specific to (**B & C**) Cyclin D1, (**E & F**) ERα, or (**H & I**) CXXC5. All experiments were repeated two independent times; representative images for WB from the same experiment are shown. Molecular masses in kDa are indicated.

Cyclin D1, which is an estrogen-responsive gene product critical for cell cycle progression through the G1/G0 to the S phase^61, 62^ and used here as a control, in E2-treated cells at the time of drug treatment (0 h) showed substantially augmented transcript and protein levels in comparison with those in untreated cells (Fig. 2A-C). ActD treatment of MCF7 cells led to a steady decline in the protein levels of Cyclin D1 levels with a half-life of about 3 h when E2 was present but it had no effect on protein levels in the absence of E2. CHX, on the other hand, rapidly and effectively repressed the protein levels of Cyclin D1 which was further augmented in the presence of ActD independently of E2 with a half-life of about 30 min, consistent with previous studies^63^. Thus, while steady-state levels of Cyclin D1 transcripts somewhat contribute to protein levels, Cyclin D1 is an unstable protein whose levels are primarily mediated post-translationally whether or not cells are treated with E2.

In contrast to Cyclin D1, ERα levels, as expected, in cells treated with E2 were markedly lower than those observed in untreated cells without substantial alterations in transcript levels (Fig. 2D-F). ERα protein levels showed a peculiar pattern in the presence of ActD: the reduced protein levels of ERα in the presence of E2 steadily accumulated reaching levels comparable to those in the absence of E2, despite the similar ERα transcript levels in the absence or presence of ActD and/or CHX independently of E2 (Fig. 2D). This suggests that a pool of ERα transcripts with delayed processing, as reported previously^64^, and/or stored/segregated in an intracellular sub-compartment(s) re-enters translation. On the other hand, treating cells with CHX alone and/or with ActD induced a progressive decline in ERα protein levels with a half-life of about 8 h independent of E2. These observations imply that pre- and post-translational mechanisms primarily mediate the levels of the ERα protein in MCF7 cells.

Although E2 treatment augmented transcript levels of CXXC5, the pattern of reduction in CXXC5 transcript levels did not differ among cells treated with ActD and/or CHX independently of E2 (Fig. 2G). E2 treatment also enhanced CXXC5 protein levels (Fig. 2H & 2I) observed similarly for Cyclin D1. Protein levels of CXXC5 showed a faster decline in the absence of E2 with a half-life of 2 h as opposed to 6 h when cells were treated with E2 (Fig. 2H & 2I). This suggests that E2 treatment of cells contributes to the stability of the CXXC5 protein. Thus, E2-mediated modulation of the CXXC5 protein levels in cells occurs through post-transcriptional and post-translational processes.

### Degradation of CXXC5 by the ubiquitin-proteasome system (UPS)

The UPS is responsible for the degradation of the majority of intracellular proteins as a multi-enzyme process encompassing the covalent conjugation of ubiquitin to substrate proteins and their recognition and degradation by the proteasome^65, 66^. To initially examine whether or not the regulation of stability of CXXC5 protein involves the UPS, we used MCF7 cells synchronized in G0/G1 with CD-FBS for 72 h and treated without or with 10^-9^ E2 for 36 h. Cells were then incubated in a fresh medium without (DMSO as the vehicle) or with a proteasome inhibitor MG132 at 10 µM concentration, which was based on preliminary studies (Supplementary Information, Fig. S3), for 4h (Fig. 3). Collected cells were then processed for and subjected to WB. Similar to ERα, levels of CXXC5 showed an increase in the presence of MG132 (+ MG132) in cells treated without or with E2. This was correlated with the increased detection of ubiquitinated cellular proteins by a ubiquitin antibody in WB (Fig. 3A & 3B). This together with our observation that bafilomycin A1 (BAF), which is an inhibitor of lysosome activity^67, 68^ did not restore the reduced levels of CXXC5 but effectively increased levels of the p62 protein, also known as autophagosome cargo protein sequestosome 1 (SQSTM1), a ubiquitin-binding scaffold protein degraded by lysosome^69, 70^ suggests that CXXC5 is degraded by the UPS (Fig. 3C). To further assess the degradation of CXXC5 by the UPS, we examined the protein stability of CXXC5 in the absence or presence of MG132. We synchronized MCF7 cells at G0/G1 with the hormone withdrawal and treated cells without or with 10^-9^ M E2 for 36 h. Cells were then incubated in a fresh medium without (DMSO as the vehicle) or with 10 µM MG132 together with 50 µg/ml CHX to inhibit translation in the absence (- E2) or presence of 10^-9^ M E2 (+ E2) for 0, 0.5, 1, 2, 4, and 8 h. The pattern of decline was similar when protein synthesis was blocked (Fig. 3D-H). MG132 effectively blocked time-dependent decreases in CXXC5 levels whether or not the cells were treated with E2. Thus, CXXC5 is degraded by the UPS.

**Fig. 3.**
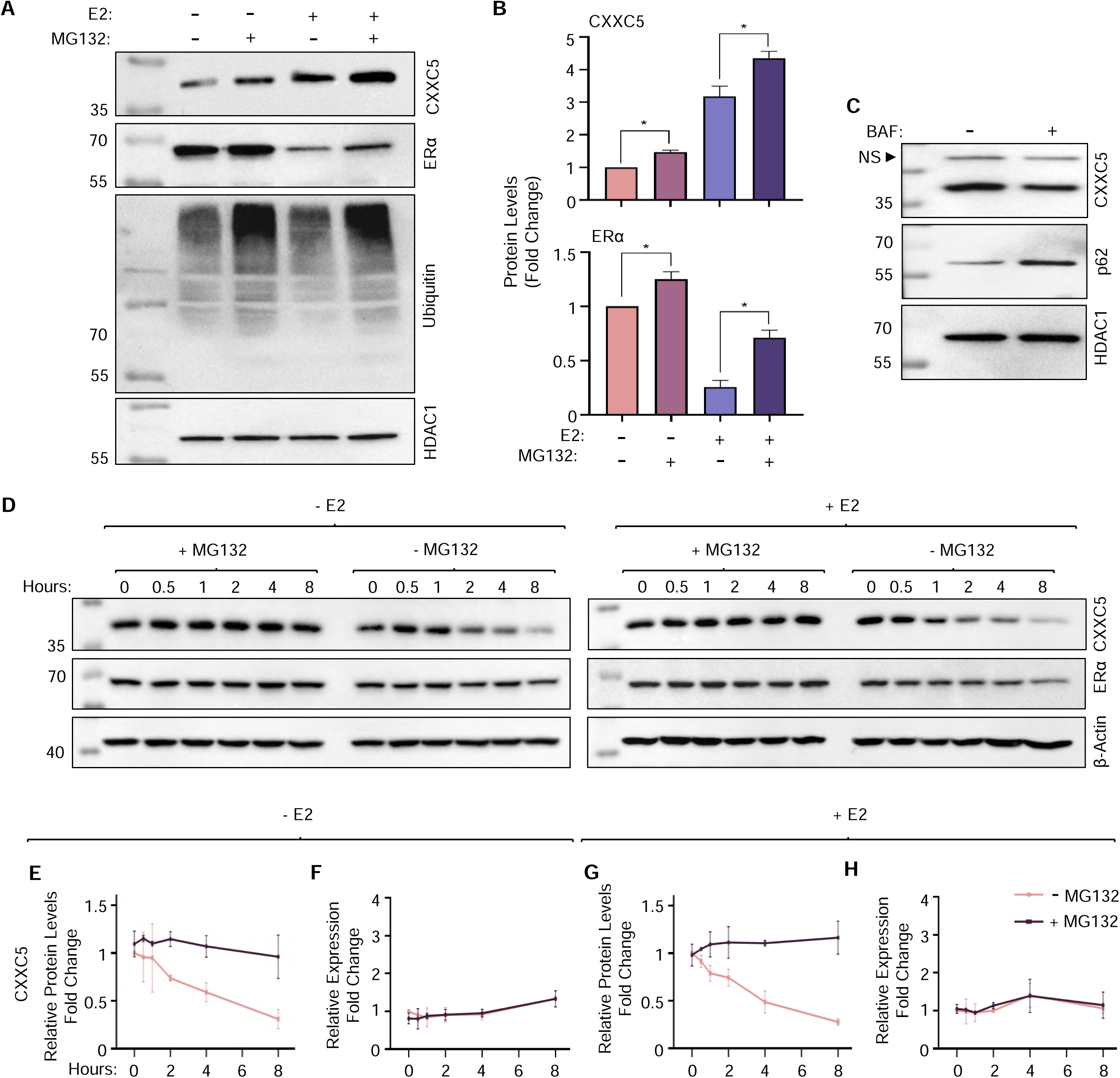
CXXC5 is degraded through the ubiquitin-proteasome system (UPS). MCF7 cells synchronized in G0/G1 with CD-FBS for 72h were treated without (- E2) or with 10^-9^ M E2 (+ E2) for 36 h. Cells were then incubated in a fresh medium without (DMSO as the vehicle) or with 10 µM MG132, a proteasome inhibitor for 4 h in the absence (- E2) or presence of 10^-9^ M E2 (+ E2). (**A**) Collected cells were processed for and subjected to WB using an antibody specific for CXXC5, ERα as control, ubiquitin, or HDAC1. Molecular masses in kDa are indicated. (**B**) Densitometric analyses of protein levels relative to that of HDAC1 in three independent determinations are indicated in the graphs. Asterisks indicate significant differences when MG132 is present. (**C**) To assess the specificity of CXXC5 degradation by the UPS, MCF7 cells were incubated in a fresh medium with a lysosome inhibitor bafilomycin A1 (BAF, 100 nM) for 4 h. Cells extracts were processed for and subjected to WB using antibodies specific to CXXC5, p62/SQSTM1 or HDAC1. NS indicates a non-specific protein. (**D-H**) Synchronized MCF7 cells for 72 h were treated without (- E2) or with 10^-9^ M E2 (+ E2) for 36 h. Cells were then treated in the absence or presence of E2 without or with 50 µg/ml CHX and 10 µM MG132 for 0, 0.5, 1, 2, 4, and 8 h. Cells were then collected for RNA and protein isolation. (**D**) Cellular extracts were subjected to WB using an antibody specific for CXXC5, ERα, or β-Actin. β-Actin was used as the loading control. Molecular masses in kDa are shown. (**E & G**) Densitometric analyses of CXXC5 protein levels relative to β-Actin levels in two independent determinations are indicated in the graphs. (**F & H**) RNAs were subjected to RT-qPCR for CXXC5 and RPLP0 mRNA levels and CXXC5 levels were normalized to *RPLP0* transcripts as a control gene.

Since CXXC5 protein levels reached the highest levels within 6 h of E2 treatment corresponding to the G0/G1 phase of synchronized MCF7 cells and remained at similar levels in subsequent phases, we asked whether or not the degradation of CXXC5 by UPS involves a specific cell cycle phase. To explore this issue, we used a cycle phase enrichment approach of synchronized MCF7 cells (Fig. 4). To examine the CXXC5 degradation by UPS at G0/G1, we treated the G0/G1 enriched cell population following 72 h of hormone withdrawal without (DMSO, 0.01% as vehicle control) or with MG132 for 4 h. We treated synchronized cells with 10^-9^ M E2 for 21 h to obtain the S-phase enriched cell population. We then incubated cells for an additional 4 h without or with MG132 in the presence of E2. Nocodazole is an anti-mitotic agent that reversibly interferes with the polymerization of microtubules by binding to β-tubulin thereby impairing the formation of the metaphase spindles during the cell division cycle^71^. To enrich the cell population in the G2/M phase, synchronized cells treated with E2 for 21 h were incubated in fresh medium containing for 2 h in the presence of E2 and 0.3 µM nocodazole, the concentration of which was based on our previous observation (submitted for publication). Cells were then treated without or with MG132 in the presence of E2 and nocodazole for an additional 4 h. Cells were collected, processed, and subjected to flow cytometry (Fig. 4A) and WB (Fig. 4B).

**Fig. 4.**
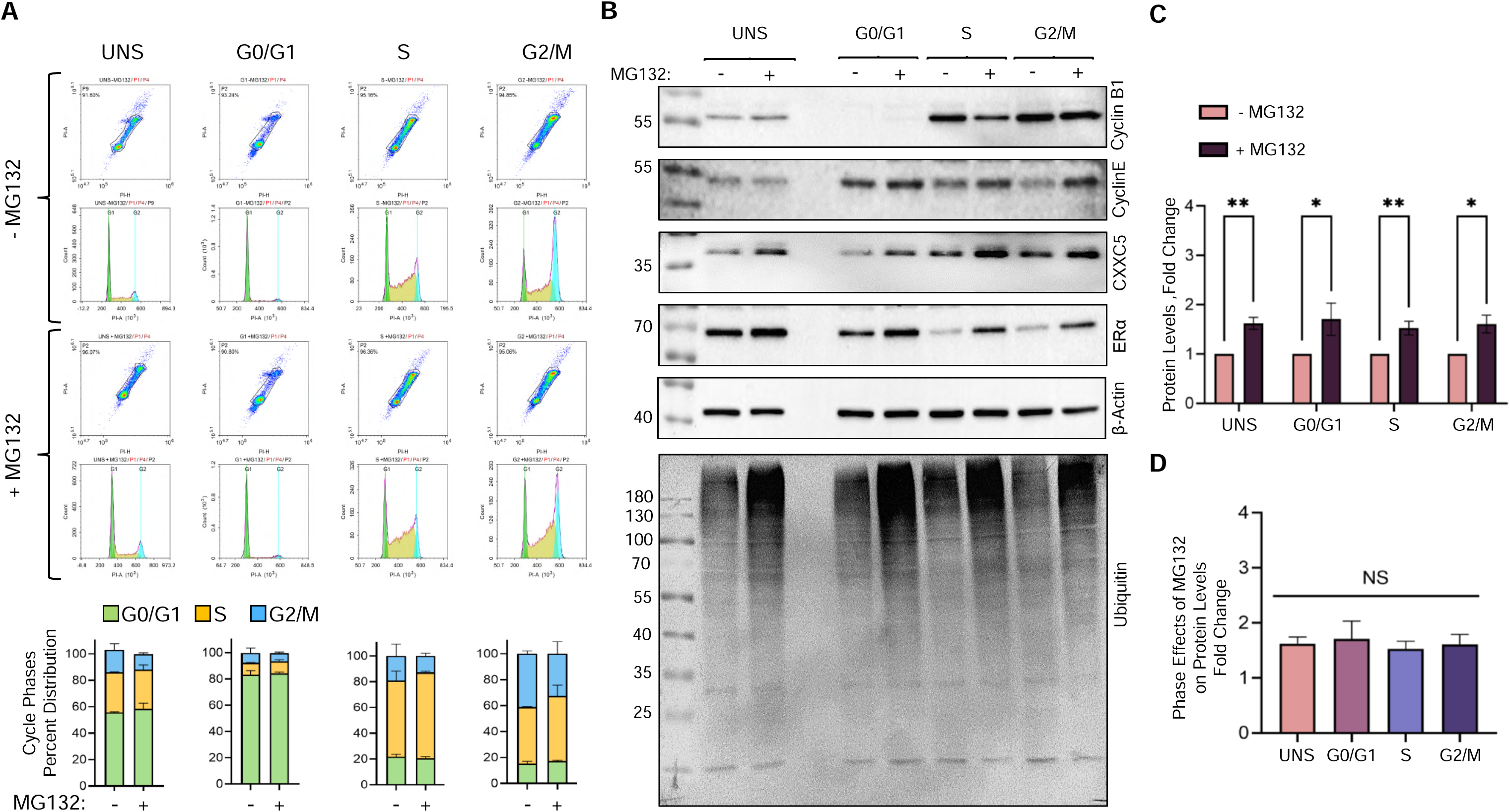
Degradation of CXXC5 by UPS at cell cycle phases of MCF7 cells. MCF7 cells synchronized at the G0/G1 phase for 72 h were treated without (DMSO, 0.01% as vehicle control) or with MG132 for 4 h. Similarly, cells synchronized at G0/G1 were incubated in a fresh media containing CD-FBS without or with 10^-9^ M E2 for 21 h to enrich the cell population at the S phase. Cells were then incubated for an additional 6 h without or with MG132 in the presence of E2. To enrich the G2/M phase we used nocodazole. For this, synchronized cells treated with E2 for 21 h were maintained for 2 h in the presence of E2 and 0.3 µM nocodazole, then another 4 h in the presence of E2 and 0.3 µM nocodazole without or with MG132. Collected cells were processed for and subjected to (**A**) flow cytometry for the assessment of cell cycle phase distributions also depicted as bar graphs, and (**B**) WB analyses using an antibody specific for Cyclin B1, Cyclin E, CXXC5, ERα, β-Actin, or ubiquitin. (**C & D**) Densitometric analyses of CXXC5 protein levels relative to those of β-Actin in three independent determinations are indicated in the graphs. Asterisks designate significant differences when MG132 is present. Molecular masses in kDa are indicated.

With expected phase distributions following E2 treatment of synchronized MCF7 cells, MG132 treatment of cell populations enriched at G0/G1, S, and G2/M phases led to an increase in the detection of ubiquitinated cellular proteins with WB using a ubiquitin-specific antibody, indicating compromised proteasome functions. We observed that levels of Cyclin B1 decreased when cells were treated with MG132 in the S phase, likely due to the repression of Cyclin B1 translation^72^. We also observed that when nocodazole-treated cells to block M phase exit were also treated with MG132 Cyclin B1 levels remained unaltered, consistent with the importance of proteolysis in mitotic exit^73, 74^. On the other hand, MG132 treatment prevented the degradation of Cyclin E, ERα, and CXXC5 without an effect on β-Actin levels used as control (Fig. 4B). Furthermore, changes in CXXC5 levels (Fig. 4C) as assessed by the ratio of protein levels in the absence and presence of MG132 remains similar in cell cycle phases (Fig. 4D). This suggests that the UPS-mediated degradation of CXXC5 is independent of cell cycle phases.

### Ubiquitination of CXXC5

Degradation of CXXC5 by UPS independent of cell cycle phases implies the ubiquitination of CXXC5 in MCF7 cells. The ubiquitination encompasses a three-step enzymatic cascade involving E1, E2, and E3 enzymes that results in the transfer of ubiquitin to a lysine (**K**) residue on substrates as well as the removal of ubiquitin modifications from substrates by de-ubiquitinase enzymes, DUBs^75, 76^. Due to dynamic and spatiotemporal regulation of protein ubiquitination^77, 78^ together with a relatively low abundance of CXXC5 protein in MCF7 cells, our several attempts to identify endogenously ubiquitinated CXXC5 protein with various approaches yielded no discernable results (Fig. 5A & 5B). To circumvent this issue, we used the ^bio^Ub approach, which is based on the use of cellular biotinylation of ubiquitin that allows the efficient isolation and enrichment of ubiquitinated proteins due to the strength and the specificity of the avidin-biotin interaction^46, 47^. The ^bio^Ub approach exploits the use of engineered six tandem ubiquitin (Ub_6_) peptides genetically fused to the BirA enzyme^46, 47^. Each ubiquitin peptide in the Ub_6_-BirA proprotein contains the BirA recognition sequence with a single biotin acceptor **K** (MGLNDIFEAQ**K**IEWHE) at the amino terminus^46, 47^. Upon synthesis, digestion of the Ub_6_-BirA protein by endogenous DUBs generates single ubiquitin peptides for biotinylation by the BirA, ^bio^Ub. Endogenous ubiquitin-conjugating enzymes then incorporate the generated ^bio^Ub into protein substrates. To explore the labeling of CXXC5 with ^bio^Ub we used HEK293FT cells, which are E2-nonresponsive and synthesize CXXC5 at low levels (Supplementary Information Fig. S4), and show high efficiencies for transfection and heterologous protein synthesis^79^. We transiently transfected HEK293FT cells with the pCAG-(Ub)_6_-BirA or the pCAG-BirA expression vector^47^ together with the pcDNA3.1(-) expression vector bearing a CXXC5 cDNA with sequences that encode amino-terminally located 3xFlag epitope, 3F,^80^ in the growth medium containing 50 µM biotin and 1 mM ATP for 24 h. Cellular extracts were subjected to precipitation using NeutrAvidin-conjugated agarose beads followed by WB using the Flag, ubiquitin, or biotin antibody. We observed an enrichment of 3F-CXXC5 with distinct molecular masses varying from 40 to 130 kDa with a predominant CXXC5 species of about 55 kDa by the use of the Flag antibody only in the NeutrAvidin precipitated ^bio^Ub_6_-BirA group which also contains highly abundant ^bio^ubiquitinated cellular proteins assessed with reprobing the membrane with the ubiquitin and biotin antibodies (Fig. 5C-E). As observed in MCF7 cells (Fig. 5A & 5B), we are uncertain about the identity of the protein species migrating at 40 kDa (indicated by a question mark in Fig. 5C). This could represent CXXC5 bound to beads, a non-specific protein bound to the beads with electrophoretic mobility similar to CXXC5, or a degraded ubiquitinated form of CXXC5. However, the latter is unlikely, as we see this protein species across all precipitated CXXC5 variants, including the variant lacking lysine residues (K_Null_, Fig. 9). Nonetheless, these findings suggest that CXXC5 is ubiquitinated in HEK293FT cells.

**Fig. 5.**
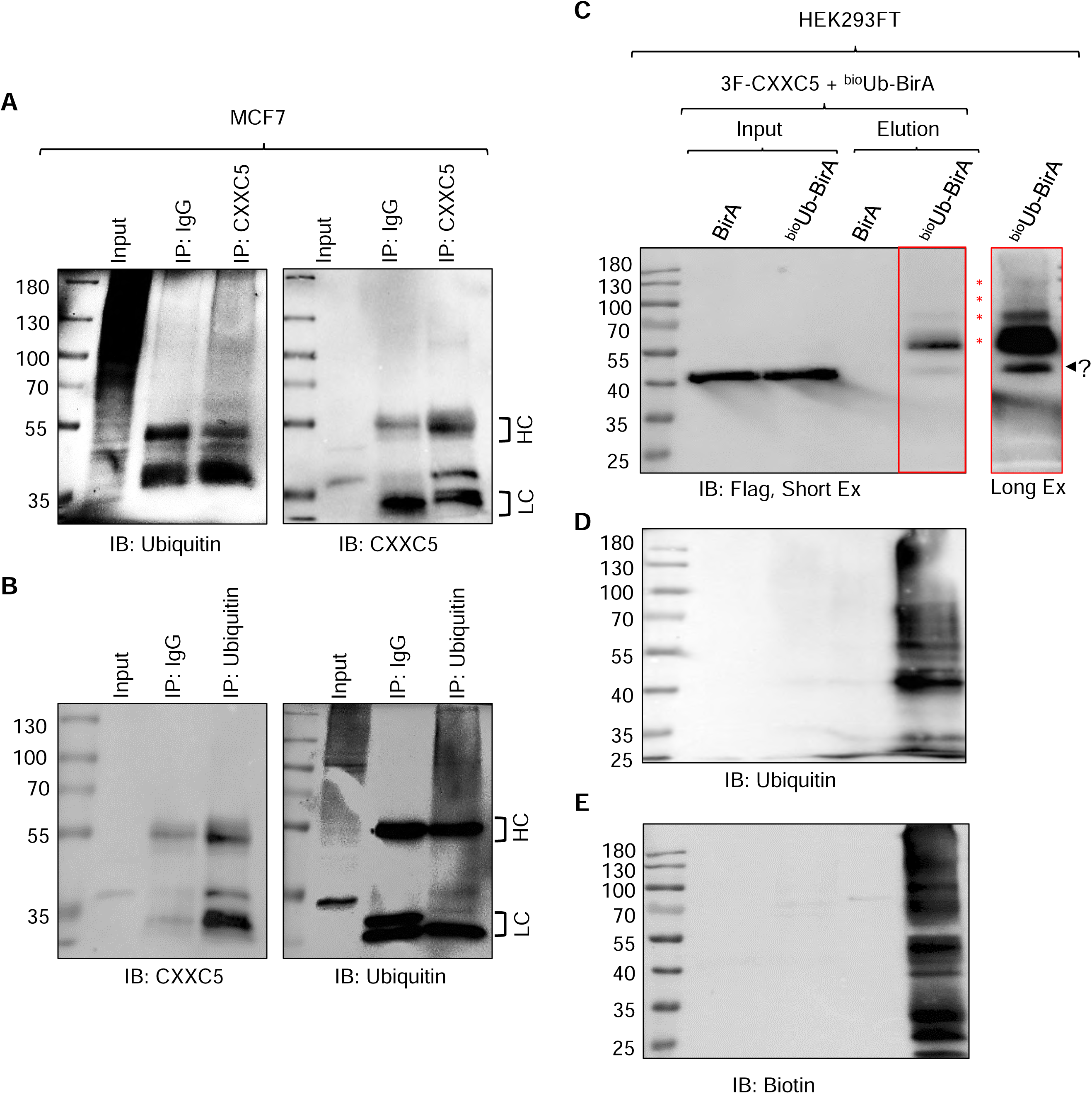
Assessing the ubiquitination of CXXC5. **(A & B)** Based on the findings that the UPS-mediated degradation of CXXC5 is independent of cell cycle phases, we carried out immunoprecipitations using an antibody specific to CXXC5 or ubiquitin with extracts of unsynchronized MCF7 cells grown in steady-state conditions followed by WB analyses using the ubiquitin, the CXXC5 antibody or isotype-matched IgG. Input indicates 5% cellular extracts used in immunoprecipitations. HC and LC denote heavy and light chains, respectively. Molecular masses in kDa are indicated. The arrow with a question mark indicates an unknown protein species. (**C-E**) Ubiquitination of CXXC5 in HEK293FT cells transiently transfected with an expression vector bearing 3F-CXXC5, BirA, and/or Ub_6_-BirA cDNA for 24 h. Cellular extracts were subjected to immunoprecipitation using NeutrAvidin-conjugated agarose beads. Membranes were probed with the (**C**) Flag, (**D**) Ubiquitin, or (**E**) Biotin antibody. Input indicates 5% cellular extracts used in immunoprecipitations. Red stars indicate possible ubiquitinated CXXC5 species. The arrow with a question mark indicates an unknown protein species.

Based on this finding, we attempted to identify ubiquitinated residue(s) in CXXC5. For this, we carried out large-scale sequential immunoprecipitations (IPs) using the Flag antibody fragment-conjugated beads followed by the ^bio^Ub pull-down using NeutrAvidin-conjugated agarose beads of cell extracts transfected with expression vectors bearing 3F-CXXC5 and (Ub)_6_-BirA cDNAs to enrich the ^bio^Ubi-CXXC5. Samples were then subjected to SDS-10% PAGE, stained with Coomassie blue, and stained protein bands on the gel were excised. Samples were in-gel digested with trypsin and the generated peptides were subjected to Liquid Chromatography-Tandem Mass Spectrometry (LC-MS/MS) to detect peptide remnants derived from ubiquitin.

Trypsin cleaves specifically peptide bonds at the carboxyl side of K and arginine (R) residues except for when K or R is followed by proline, P^81^. The carboxyl-terminus of the mature ubiquitin terminates with ^71^LRLRGG^76^ amino acid sequence in which the last Gly^76^ residue is conjugated to primarily K residues^82, 83^ on target proteins^84, 85^. In addition to target proteins, the peptide bonds at the carboxyl side of R^74^ at the carboxyl-terminus of ubiquitin when conjugated to target proteins or in free form are also cleaved by trypsin. This cleavage leaves two glycine residues, G^75^G^76^, on the conjugated lysine residues of substrates generating ‘ubiquitin signature peptides’ ^86, 87^ with a monoisotopic mass of 114.04 Da on the modification sites. Although rare, trypsin also cleaves the peptide bond between the penultimate R^72^ and leu^73^ (L) residues of the carboxyl-terminus of ubiquitin conjugated to a substrate. This generates an LRGG peptide with a monoisotopic mass of 383.23 Da^86, 87^. These unique K-conjugated ubiquitin remnants are used to identify the ubiquitination sites through mass spectrometry. It should be noted that the G^75^G^76^ remnant on ubiquitinated peptides after tryptic digestion is not unique to ubiquitin, but is shared with NEDD8 and ISG15, members of the ubiquitin-like protein family, UBL^88^. This necessitates using ubiquitination-specific approaches including ^bio^ubiquination as we utilized here for identifying ubiquitinated K residue(s). CXXC5 contains ten K residues (red font in Fig. 6B) and seven R residues (blue font in Fig. 6B) at the amino terminus, and seven K and seven R residues in the CXXC domain at the carboxyl terminus of the protein. In addition, four K residues and two R residues are present in a region between the amino terminus and the CXXC domain of CXXC5 that contains the canonical monopartite nuclear localization signal (cNLS, ^258^KKKRKR^262^) ^89^. The locations of both K and R residues in CXXC5 present 34 possible trypsin cleavage sites, assessed with PeptideCutter (https://web.expasy.org/peptide_cutter/) with two exceptions: ^77^R and ^301^K which are followed by proline residues. This potentially generates many cleaved single K residues by trypsin with possible ubiquitin signatures along with peptides with lengths ranging from two to more than 30 amino acids. Since the optimal peptide length for mass spectrometric detection of peptide fragments is 8-25 amino acids^90^, the frequency and location of trypsin cleavage sites rendered the identification of ubiquitinated K residues of 3F-CXXC5 tryptic peptides difficult. Nevertheless, LC-MS/MS results revealed two tryptic CXXC5 peptides containing K residues that bear ubiquitin signatures: K^147^ in ^138^SGAVASLLS**K**AER^150^ and K^309^ in ^302^KPSAALE**K**VMLPTGAAFR^319^ peptides. Previous studies on the ubiquitome landscape of cell models reported that CXXC5 is one of the ubiquitinated proteins containing ubiquitinated K^147^ in ^138^SGAVASLLS**K**AER^150 (88, 91)^ and K^292^ in ^286^TGHQIC**K**FR^294^ peptide sequence^88^. Confirming the ubiquitination of K^147^ and identifying the unique K^309^ ubiquitination, our, together with others, results indicate that CXXC5 can be ubiquitinated at multiple K residues located in both the amino-terminus region and the carboxyl-terminus CXXC domain.

**Fig. 6.**
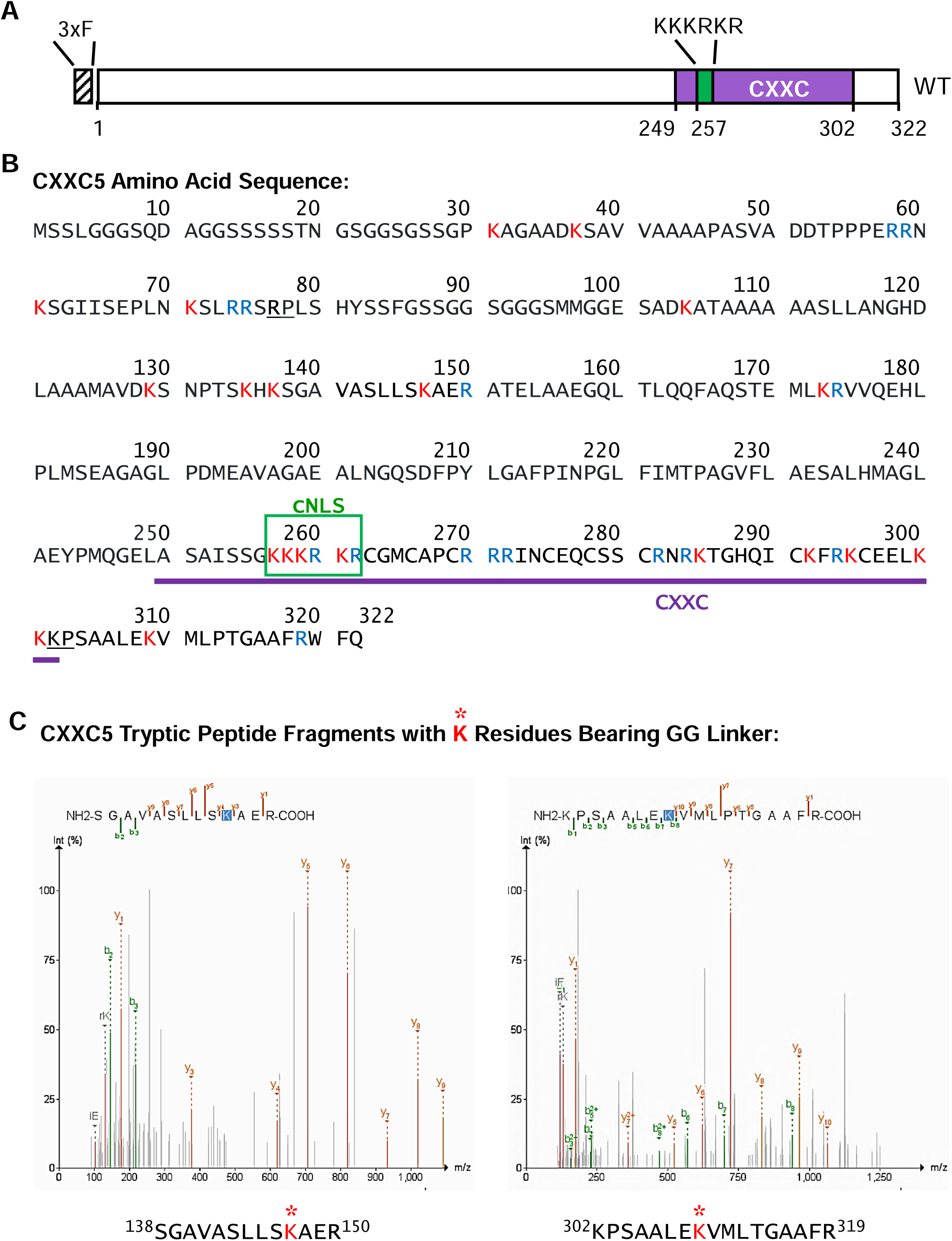
Identification of ubiquitinated lysine residues of CXXC5. (**A**) Schematics of the CXXC5 protein. 3xF (in black stripe) indicates the 3xFlag epitope, NLS (in green font and rectangle) denotes the nuclear localization signal, and CXXC (in purple font and underlines) defines the DNA binding domain. (**B**) The amino acid sequence of CXXC5. Possible lysine (K in red font) and arginine (R in blue font) residues that are targets for trypsin cleavage are indicated. (**C**) Identified peptides containing ubiquitinated lysine residue (in red font and marked with asterisks) from MS analysis. For the identification of ubiquitinated lysine residues, HEK293FT cells were transiently transfected with the pcDNA-3F-CXXC5 and with the pCAG-(Ub)_6_-BirA or the pCAG-BirA expression vector in a growth medium containing 50 µM biotin and 1 mM ATP for 24 h. Cellular extracts were subjected to sequential immunoprecipitation using the Flag antibody fragment-conjugated beads followed by the ^bio^Ub pull-down with NeutrAvidin-conjugated agarose beads. Samples were then subjected to SDS-10%PAGE, stained with Coomassie blue, and stained protein bands on the gel were excised. Samples were in-gel digested with trypsin and the generated peptides were subjected to LC-MS/MS to detect peptide adducts derived from ubiquitin.

### Verification of ubiquitination sites of CXXC5

To verify the ubiquitination of K^147^ and/or K^309^ in CXXC5 and assess the effects on protein stability, we generated CXXC5 mutants by replacing each identified positively charged K residue alone or in combination with structurally similar a positively charged R residue with site-directed mutagenesis using overlapping PCR^48, 92^ and cloned into the pcDNA3.1(-) expression vector. In these constructs, we used the HA tag (YPYDVPDYA) rather than the 3xFlag tag, 3F, (DY**K**DHDGDY**K**DHDIDY**K**DDDD**K**) bearing a CXXC5 cDNA since 3F contains four K residues (in boldface) that could potentially be ubiquitinated, as we observed in preliminary studies (Supplementary Information, Fig. S5). We transiently transfected HEK293FT cells with the expression vector bearing WT HA-CXXC5 or a K to R changed mutant HA-CXXC5 (Fig. 7A) cDNA together with the pCAG-(Ub)_6_-BirA expression vector in the growth medium containing 50 µM biotin and 1 mM ATP for 24 h. Cellular extracts of WT, R_147_, R_309_, or R_147/309_ synthesizing cells assessed with WB using the HA antibody (Fig. 7B) were also subjected to precipitation using NeutrAvidin-conjugated agarose beads followed by WB using the HA or Biotin antibody. We observed that mutations alone or in combination did not prevent the ubiquitination of the variant proteins compared with WT HA-CXXC5 (Fig. 7C). These results suggest that the remaining K residue(s) in mutant CXXC5 proteins likely undergo ubiquitination and/or become new targets for ubiquitination. To assess whether or not mutations alter the extent or pattern of stability of CXXC5 variant proteins, MCF7 cells, grown in steady-state conditions, were transiently transfected for 48 h. Cells were incubated in a fresh medium with 50 µg/ml CHX for 0, 0.5, 1, 2, 4, and 8h. Cellular extracts were subjected to WB using the CXXC5 or HA antibody (Fig. 7D). Due to the molecular mass of the HA tag at the amino-termini, CXXC5 variants display slower electrophoretic migration in WB. This provides an opportunity to comparatively assess the degradation of both the endogenous (en) CXXC5 and HA-tagged CXXC5 variants synthesized in transfected cells (tr) in WB blots, which are also verifiable with the HA antibody. Quantitative analyses of the WB results normalized to the levels of β-Actin as the loading control revealed that R_147_, R_309_, and R_147/309_ mutant HA-CXXC5 proteins show stability patterns similar to that observed with the WT CXXC5 (Fig. 7E). These results are consistent with our conclusion that the remaining K residue(s) in mutant CXXC5 proteins are, or become, targets for ubiquitination and subsequent changes in stability.

**Fig. 7.**
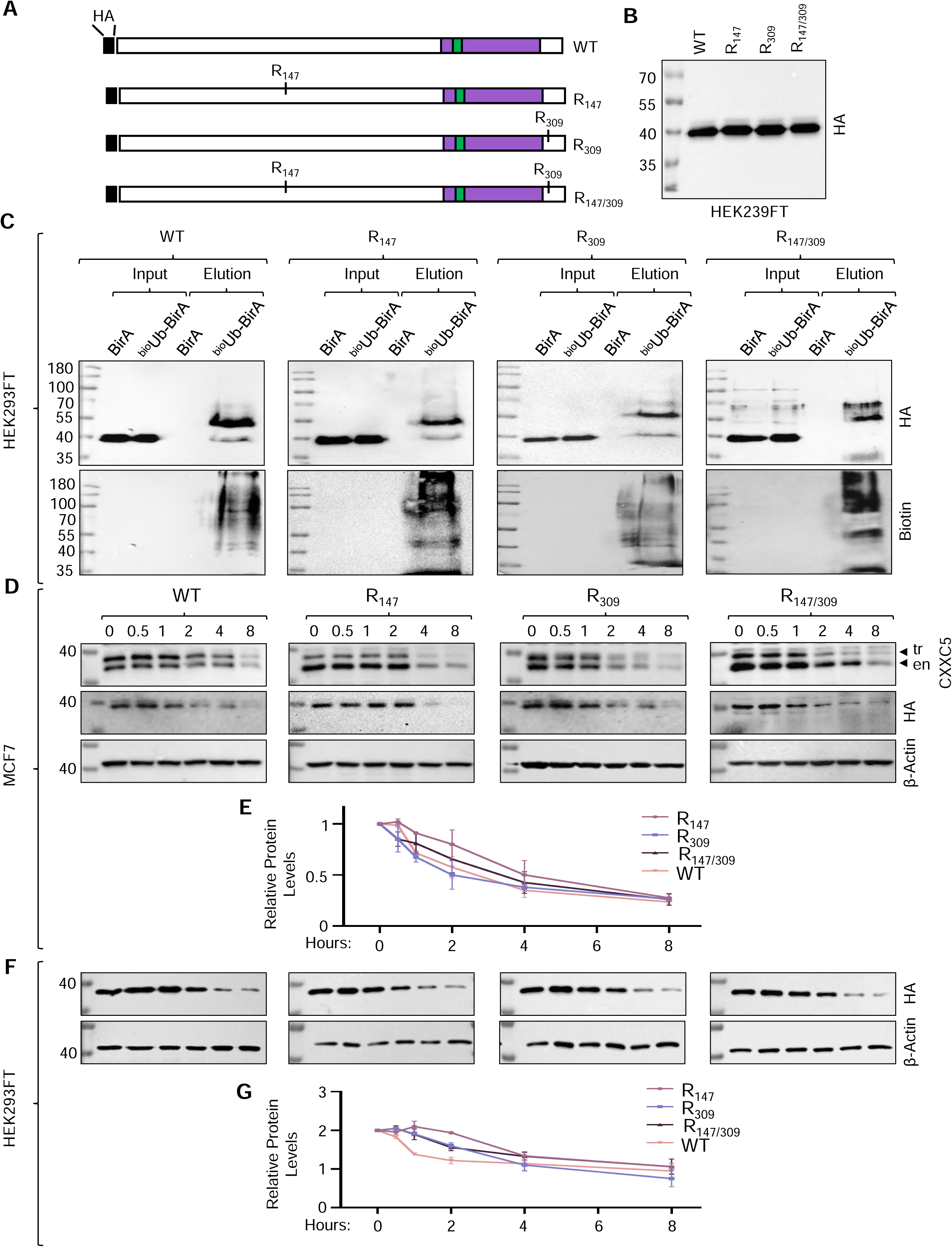
Verification of ubiquitination of K^147^ and/or K^309^ residues of CXXC5. (**A**) Schematics of mutant CXXC5 bearing R residues at the position of MS-identified ubiquitinated K^147^ and/or K^309^ residues. (**B**) HEK293FT cells were transiently transfected with an expression vector bearing either WT CXXC5 or a CXXC5 variant cDNA for 24 h. Cells were then collected for protein isolation and cellular extracts were subjected to WB using an antibody specific for HA tag to confirm the expression of mutant CXXC5 proteins. (**C**) HEK293FT cells were transiently co-transfected with an expression vector bearing a CXXC5 variant cDNA and the pCAG-(Ub)_6_-BirA or the pCAG-BirA expression vector in the growth medium containing 50 µM biotin and 1 mM ATP for 24 h. Cellular extracts were subjected to NeutrAvidin-conjugated agarose beads followed by WB using the HA antibody. Experiments were repeated two independent times; representative images for WB from the same experiment are shown. Molecular masses in kDa are indicated. (**D**) To assess the degradation of CXXC5 variants, MCF7 (**D & E**) or HEK293FT (**F & G**) cells grown in steady-state conditions were transiently transfected with an expression vector bearing a CXXC5 variant for 48 h. Cells were then treated in the absence or presence of 50 µg/ml CHX for 0, 0.5, 1, 2, 4, and 8 h. MCF7 protein extracts were subjected to WB using an antibody specific for CXXC5, HA, or β-Actin, the latter which was used as the input control. Similarly, protein extracts of transfected HEK293 cells were subjected to WB using an antibody specific for HA or β-Actin. (**E & G**) Densitometric analyses of protein levels in MCF7 and HEK293FT cells relative to β-Actin in two independent determinations as the mean with error bars are shown in the graph. Molecular masses in kDa are indicated.

Due to the number (21) of K residues in CXXC5, mutation of each K residue alone or in combination presents an overwhelming task. We reasoned that the positional conservation of the K residues observed in LC-MS/MS analysis alone or in combination in otherwise a K-null mutant CXXC5 protein (K_Null_) in which every other K residue in CXXC5 is changed to R residue would allow us to assess the ubiquitination of a specific K residue. Since CXXC5 is predominantly localized in the nucleus and the monopartite canonical NLS (cNLS) of CXXC5 contains four K residues (^258^KKKRKR^262^), we initially assessed whether or not K_Null_ bearing the canonical NLS sequence, K_Null-cNLS_, is localized to the nucleus in transiently transfected HEK293FT cells for 24 h or MCF7 cells for 48 h, at times which transgene synthesis becomes maximal (data not shown). While, as expected, K_Null-cNLS_ localized to the nucleus of both MCF7 and HEK293FT cells, K_null-cNLS_ was also ubiquitinated in HEK293FT cells as assessed with ^bio^Ub suggesting that K residues in the cNLS could undergo ubiquitination (Supplementary Information, Fig. S6). To circumvent this issue, we initially examined whether or not changing K residues in the nNLS would alter the intracellular localization of CXXC5. For this, we generated CXXC5 mutants (Fig. 8A). Of the mutants, WT_M-NLS_ bears random amino acid replacements in the cNLS changing the ^258^KKKRKR^262^ sequence to ^258^AASGGS^262^. We observe that WT_M-NLS_ largely localizes to the cytoplasm in transiently transfected HEK293FT and MCF7 cells (Fig. 8B). Besides cNLS, several proteins including capsid protein 1, VP-1^93^, contain an NLS that bears no K but R residues critical for the localization to the nucleus^94^. Based on these findings, we assessed whether ^258^RRRRRR^262^ instead of ^258^KKKRKR^262^ sequence produces a mutant CXXC5 protein that localizes to the nucleus. We transiently transfected cells with the expression vector bearing WT HA-CXXC5 (WT_R-NLS_) or the K_Null_ CXXC5 mutant with the ^258^RRRRRR^262^ NLS motif (K_Null_) cDNA. As WT_R-NLS_, K_Null_ localized to the nucleus in transiently transfected HEK293FT and MCF7 cells (Fig. 8B), suggesting that the ^258^RRRRRR^262^ sequence is a monopartite NLS motif. We further verified these results with a GFP protein that bears the ^258^RRRRRR^262^ motif at the amino-terminus, GFP_R-NLS_. While the WT GFP showed diffuse intracellular staining encompassing the cytoplasm and nucleus, GFP_R-NLS_ predominantly localizes to the nucleus (Fig. 8B).

**Fig. 8.**
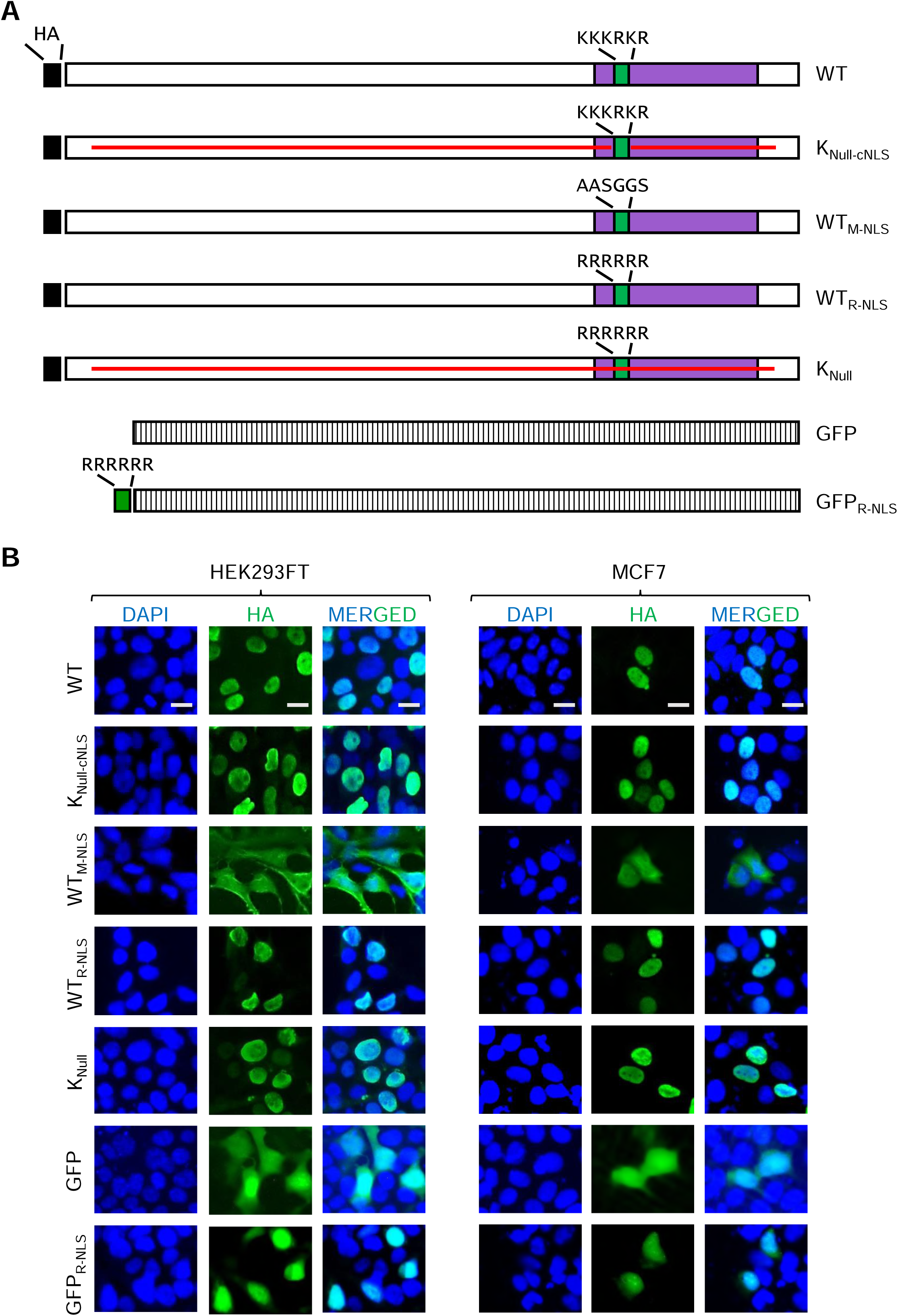
Intracellular locations of CXXC5 variants. (**A**) Schematics of HA-tagged WT and CXXC5 variants without or with mutations at the NLS. GFP indicates green fluorescent protein without or with an NLS of R residues at the amino terminus of the protein (GFP_R-NLS_). (**B**) We transiently transfected HEK293FT cells or MCF7 cells with an expression vector bearing cDNA for 24 h or 48 h respectively. Cells were then subjected to immunocytochemistry using the HA antibody followed by an Alexa Fluor 488-conjugated goat-anti-rabbit secondary antibody. DAPI was used for DNA staining. Images were captured with a fluorescent microscope. The scale bar is 10 µm.

Based on these results, we assessed the ubiquitination of HA-K_Null_ mutant CXXC5, K_Null_, in comparison with WT HA-CXXC5, WT, using the ^bio^Ub approach in transiently transfected HEK293FT cells (Fig. 9A-E). Although the estimated molecular mass of the K_Null_ mutant is comparable to that of WT, K_Null_ shows an electrophoretic migration somewhat faster than the WT in WB using the HA antibody (Fig. 9B). Precipitation of protein extracts of transiently transfected HEK293FT cells with the ^bio^Ub approach using NeutrAvidin-conjugated agarose beads followed by WB using the biotin antibody indicates that K_Null_ is not ubiquitinated in contrast to WT HA-CXXC5 (Fig. 9C). Furthermore, we find in transiently transfected MCF7 cells treated with 50 µg/ml CHX for 0, 0.5, 1, 2, 4, and 8h that K_Null_ is stable compared to WT HA-CXXC5 (tr) or endogenous CXXC5 (en) assessed with the CXXC5 or HA antibody (Fig. 9D & 9E), as we similarly observed in transiently transfected HEK293FT cells assessed with the HA antibody (Fig. 9F & 9G). These results suggest that K residues of CXXC5 are targets for ubiquitination and are critical for the stability of the protein.

**Fig. 9.**
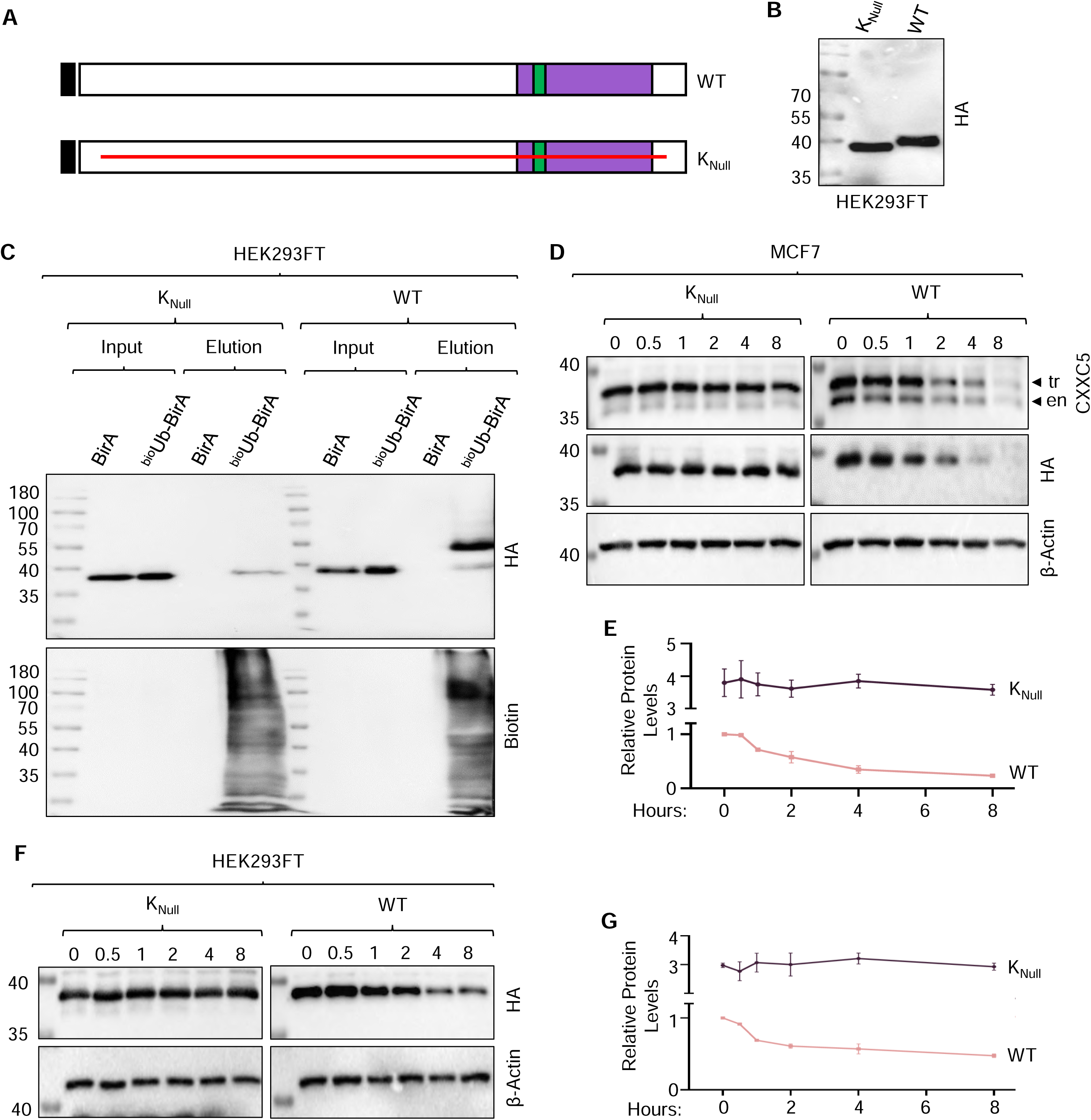
Ubiquitination and stability of the HA-Lysine-free (K_Null_) CXXC5 variant. (**A**) Schematics of the HA-tagged WT and the K_Null_ CXXC5 variant. (**B & C**) Cellular extracts of transiently transfected HEK293FT cells with an expression vector bearing a K_Null_ mutant CXXC5 variant cDNA and the pCAG-(Ub)6-BirA or the pCAG-BirA expression vector for 24 h were subjected to WB using HA antibody. Molecular masses in kDa are indicated. (**C**) Ubiquitination of K_Null_ was assessed in transiently transfected HEK293FT cells. Cellular extracts were subjected to precipitation using NeutrAvidin-conjugated agarose beads followed by WB using the HA antibody. Membranes were re-probed with the biotin antibody. Molecular masses in kDa are shown. (**D**) In assessing the stability of HA-K_Null_ in comparison with WT HA-CXXC5, protein extracts from transiently transfected MCF7 cells for 48 h and treated with 50 µg/ml CHX for 0, 0.5, 1, 2, 4, and 8 h were subjected to WB with the antibody specific to CXXC5, HA or β-Actin. All experiments were repeated two independent times; representative images for WB from the same experiment are shown. Molecular masses in kDa are indicated. (**E**) The Graph as the mean with error bars shows densitometric analyses of protein levels normalized to β-Actin in two independent determinations. (**F**) To examine the stability of K_Null_ in comparison with WT, we transiently transfected HEK293FT cells for 24 h. We then treated the cells in the presence of 50 µg/ml CHX for 0, 0.5, 1, 2, 4, and 8 h. Cellular extracts were subjected to WB using an antibody specific for CXXC5 or HA antibody. We used β-Actin levels as the loading control using the β-Actin antibody (β-Actin). All experiments were repeated two independent times; representative images for WB from the same experiment are shown. Molecular masses in kDa are indicated. (**G**) Densitometric analyses of protein levels normalized to β-Actin in two independent determinations are shown as the mean with error bars in the graph.

We then wanted to verify the ubiquitination of K_147_ and/or K_309_ by changing R^147^ and/or R^309^ residues in the K_Null_ mutant back to K residue(s) (Fig. 10A-10F). We precipitated synthesized CXXC5 variants (Fig. 10B) in cellular extracts from HEK293FT cells transiently co-transfected with expression vectors bearing ^bio^Ub and CXXC5 variant cDNAs for 24 h with NeutrAvidin-conjugated agarose beads followed by WB using the HA or Biotin antibody. Results show that mutant CXXC5 protein bearing K^147^ (K_Null&K147_), K^309^ (K_Null&K309_), and K^147/309^ (K_Null&K147/309_) indeed undergo ubiquitination (Fig. 10C). To examine the effects of ubiquitination on the stability of these mutant CXXC5 proteins, we transiently transfected MCF7 cells, grown in steady-state conditions, for 48 h. Cells were then incubated in a fresh medium with 50 µg/ml CHX for 0, 0.5, 1, 2, 4, and 8h. Cellular extracts were subjected to WB using the CXXC5 or HA antibody. We utilized protein levels of β-Actin for the loading control in WB using the β-Actin antibody (Fig. 10D). Quantitative analysis of WBs indicates that K_Null&K147_ exhibits a degradation pattern similar to those of endogenous CXXC5. However, we observed little to no synthesis of K_309_ and K_Null&K147/309_ in WBs (Fig. 10D) whether or not the cells were treated with MG132 for 8 h (Supplementary Information, Fig. S7). As in MCF7 cells, K_Null&K147_ was similarly degraded in HEK293FT cells, whereas K_Null&K309_ was notably stable (Fig. 10E & 10F). In contrast, K_Null&K147/309_ is degraded more slowly than K_Null&K147_ (Fig. 10E & 10F). These findings suggest that ubiquitination at K_147_ is essential for CXXC5 degradation, whereas K_309_ alone does not influence degradation but mitigates the impact of K_147_ on CXXC5 degradation in HEK293FT cells. The failure of detection of K_Null&K309_ and K_Null&K147/309_ even in simultaneous transient transfections in MCF7 cells under the same conditions with identical reagents carried out with HEK293FT cells suggests that K_Null&K309_ or K_Null&K147/309_ are insufficiently translated or undergo co-translational degradation in MCF7 cells. These findings suggest that the ubiquitination of specific K residue(s) involved in CXXC5 degradation is cell-type-dependent.

**Fig. 10.**
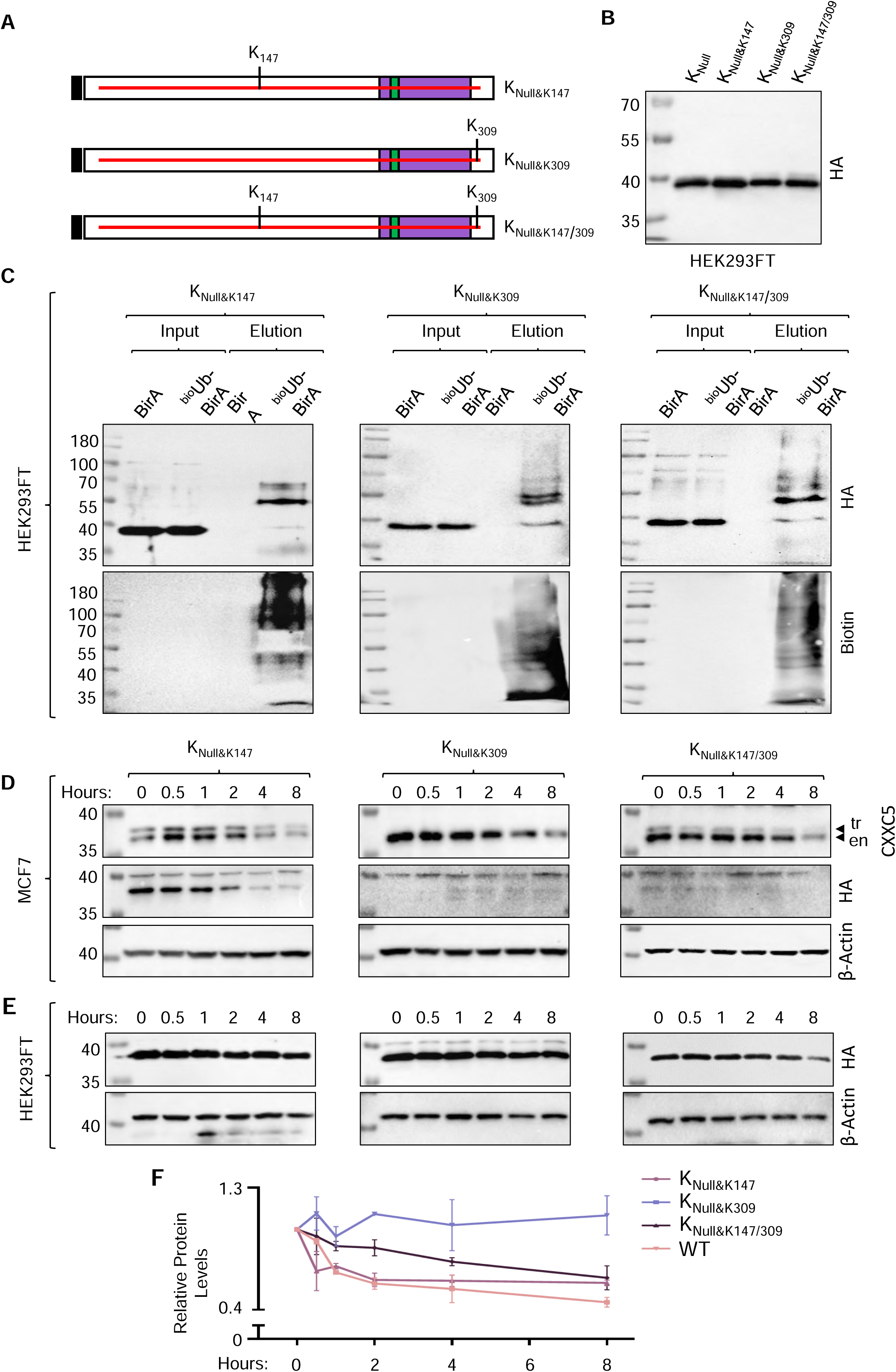
Ubiquitination and stability of the K_Null_ CXXC5 mutants. (**A**) Schematics of the K_Null_ mutant CXXC5 variants with K residues identified with MS as ubiquitinated. (**B**) Protein extracts of transiently transfected HEK293FT cells for 24 h with expression vectors bearing HA-K_Null&K147_, HA-K_Null&K309_, HA-K_Null&K147/309_, or HA-WT CXXC5 cDNA were subjected to WB using the HA antibody to assess the synthesis of variants. Molecular masses in kDa are indicated. (**C**) To evaluate the ubiquitination of variants transiently transfected HEK293FT cells for 24 h were subjected to the ^bio^Ubi approach and cellular extracts were precipitated with NeutrAvidin-conjugated agarose beads followed by WB using the HA or Biotin antibody. (**D**) The stability of mutant proteins was also assessed in transiently transfected MCF7 cells for 48 h. Cells were then treated with 50 µg/ml CHX for 0, 0.5, 1, 2, 4, and 8 h and cellular extracts were subjected to WB with the antibody specific to CXXC5, HA, or β-Actin. All experiments were repeated two independent times; representative images for WB from the same experiment are shown. Molecular masses in kDa are indicated. (**E**) To examine the stability of mutant proteins, we also transiently transfected HEK293 cells for 24 h. Cells were then incubated in the growth medium containing 50 µg/ml CHX for 0, 0.5, 1, 2, 4, and 8 h. Cellular extracts were subjected to WB using the HA antibody. β-Actin levels were used as the loading control. All experiments were repeated two independent times; representative images for WB from the same experiment are shown. Molecular masses in kDa are indicated. (**F**) Densitometric analyses of protein levels normalized to β-Actin in two independent determinations are depicted as the mean with error bars in the graph.

## DISCUSSION

Our results using the ^bio^Ub approach followed by sequential IP coupled MS analyses indicate that CXXC5 is a ubiquitinated protein and the ubiquitination contributes to the stability and degradation of the protein by UPS.

Our observations that despite a steady increase in CXXC5 transcript levels in response to E2 throughout cell cycle phases the augmented CXXC5 protein levels reach a plateau in the early G1 phase suggest a post-transcriptional regulation which critically contributes to protein levels of CXXC5. It is well established that dynamic and integrated multistep processes modulate the abundance of proteins. Although transcription, splicing, polyadenylation, export, storage, and stability of transcripts as well as cis-regulatory elements within transcripts and trans-acting factors interacting with cis-elements, including ribosomes, are critical contributors to protein abundance, the dynamics of degradation in response to various physiological conditions ultimately determines protein levels in cells^95, 96^. Referred to as translational buffering or translational offsetting, emerging evidence suggests that amounts of a set of translated mRNAs and corresponding protein levels can be maintained at constant levels despite changes in mRNA abundances ^97, 98^ due to distinct features of transcript cis-elements including 5′ untranslated regions and activity of modified tRNAs as trans-factors^98^. The structural and functional characteristics of CXXC5 transcripts remain unclear. However, specific features of CXXC5 transcripts could enable rapid and efficient protein synthesis in response to E2. This could involve a selective entrance of available CXXC5 mRNAs into the translational process and/or a selective increase in the translation efficiency of the transcripts. It is also possible that the release of stored CXXC5 transcripts from cellular compartments, such as membrane-less cytoplasmic ribonucleoprotein (RNP) assemblies including processing bodies (PBs), upon the presence of E2 may facilitate CXXC5 protein synthesis independently of transcription. This possible mechanism may also explain our observation that, when transcription is blocked, the reduced ERα protein levels in response to E2 gradually accumulate eventually reaching levels similar to those seen in the absence of E2.

Our findings that the rapid rise in CXXC5 levels within 6 h in response to E2 corresponding to the G1 phase levels off in subsequent phases despite increases in CXXC5 transcript levels also imply post-translational control including the ubiquitination of the protein, as we show here. Ubiquitination is a dynamic and spatiotemporally modulated co- and post-translational modification that regulates cellular processes through proteolytic and nonproteolytic mechanisms, including proteasomal degradation and proteostasis, selective autophagy, cell signaling cascades, protein trafficking, DNA repair and genome integrity, cell cycle control, and programmed cell death. Conjugation of ubiquitin as a monomer to one K residue on substrate proteins generates monoubiquitination; whereas, multi-monoubiquitination is the attachment of a single ubiquitin to multiple K residues on a substrate protein. Conversely, the attachment of ubiquitin molecules in a linear fashion to a single lysine (K) residue on a substrate protein results in the formation of a polyubiquitin chain. Due to the seven K residues of ubiquitin, polyubiquitination can generate linear or branched chains with different topologies. Polyubiquitination primarily marks proteins for post-translational degradation through UPS, mono- and multi-monoubiquitinated proteins can also be degraded by proteasomes^99^. Proteins are also degraded co-translationally. Co-translational ubiquitination is a robust process and a substantial fraction of polyribosome-associated nascent polypeptides are ubiquitinated during translation as a reflection of a protein quality control system that monitors protein folding during translation. This can trigger the initiation of degradation of a protein before its synthesis is complete^100, 101^.

Our LC-MS/MS results indicate that K^147^ and K^309^ of CXXC5 are ubiquitinated. The possible cell-type specific K ubiquitination is notwithstanding, previous studies on the ubiquitome landscape of cell models reported that in addition to K^147 (88, 91)^, the K^292^ residue of CXXC5 is also ubiquitinated^88^. These findings collectively suggest that CXXC5 can undergo ubiquitination at K^147^, K^292^, and K^309^ residues. However, the extent of CXXC5 ubiquitination at K residues by current approaches is likely an underestimation. This is, as we discussed in the Results section, due to the number of 1) K residues in CXXC5; 2) potential cleavage sites by trypsin due to the locations of K and R residues amenable to the enzyme; 3) generated single K residues upon trypsin digestion with possible ubiquitin signatures along with generated peptides sizes ranging from two to more than 30 amino acids in CXXC5. These factors likely contributed to the identification of ubiquitinated K residues of CXXC5 challenging through LC/MS-MS, which is most effective for detecting peptide fragments ranging from 8 to 25 residues^90^. Nevertheless, we also find here that K^147^ and/or K^309^ residues of CXXC5 are ubiquitinated. Although the type or nature of ubiquitination is unclear, these ubiquitinated K residues are involved in the stability of the protein in a manner specific to cell type. We observed that the K_Null_ mutant bearing K^147^, K_Null&K147,_ is ubiquitinated and undergoes degradation by UPS similar to the WT CXXC5 in both HEK293FT and MCF7 cells. On the other hand, the mutant CXXC5 proteins bearing K^309^ alone, K_Null&K309_, or together with K^147^, K_Null&K147/309_ that are ubiquitinated are rapidly degraded in MCF7 cells in contrast to HEK293FT cells wherein both mutant proteins are relatively stable compared to the WT CXXC5. Although the mechanism(s) is unclear, the folding of carboxy-terminally located K^309^ could differ dramatically from other K_Null_ variants leading to the recognition by the protein quality control system of MCF7 cells that initiates ubiquitination and rapid degradation of the variant. Whatever the mechanism might be, our findings indicate that the ubiquitination of specific lysine residue(s) on CXXC5 may serve as a key factor in regulating the protein’s rapid degradation when it is not needed or when its levels surpass the physiological threshold necessary for proper function in MCF7 cells.

Previously we suggested that E2-responsive CXXC5 binds to unmethylated CpG dinucleotide and acts as a nucleation factor in modulating target gene expressions critical for E2-mediated cellular proliferation in cell models synthesizing ERα^45, 48, 80^. Gene expression relies on transcription factors that bind to specific sequences in gene promoters or enhancers. Insightful kinetic studies using a well-characterized E2-responsive *TFF1/pS2* promoter indicate that the cyclic engagement of ERα with DNA directs and achieves the sequential and combinatorial assembly of transcriptional complexes including ubiquitin ligases and regulatory components of the proteasome on the gene promoter^53, 102, 103^. Subsequent ubiquitination and proteasome-mediated turnover of ERα, as well as associated co-factors, contribute to the duration and magnitude of transcriptional output^53, 102, 103^. It is therefore plausible that the stability of CXXC5 levels, which increase in response to E2 during early G1, is primarily controlled by ubiquitination when CXXC5 is bound to DNA. This regulation may occur through enhanced/decreased interactions between CXXC5 and co-regulatory proteins, ensuring the initiation of transcription and subsequently promoting its dissociation from DNA and subsequent degradation through the UPS. This is likely essential for the fine-tuning of target gene expression during cell cycle transitions, ultimately influencing cell proliferation.

In summary, we find that although E2 enhances both the transcription and synthesis of CXXC5 during the G1 phase in synchronized MCF7 cells, CXXC5 levels are primarily regulated by ubiquitination, independent of cell cycle phases. We report that multiple K residues of CXXC5 can be ubiquitinated and ubiquitinated K residues facilitate the degradation of CXXC5 via the ubiquitin-proteasome pathway. A better understanding of the interdependency of ubiquitination of and gene expressions mediated by CXXC5 could provide novel avenues for the development of new therapeutic approaches in E2 responsive ERα synthesizing target tissues including the breast wherein CXXC5 critically contributes to E2-driven cellular growth.

## MATERIALS AND METHODS

### Reagents

17β-estradiol (E2; Cat # E2257), and propidium iodide (Cat # P4170) were purchased from Sigma-Aldrich Inc. (MO, USA). Nocodazole (Cat # 31430-18-9) was obtained from Merck (Darmstadt, Germany). Antibodies for ERα (Cat # sc-543), Cyclin B1 (Cat # sc-245), Cyclin E (Cat # sc-247), and HDAC1 (Cat # sc-81598) were obtained from Santa Cruz Biotechnology (SCBT, Santa Cruz, CA, USA). The antibody for the Flag tag was from Sigma-Aldrich Inc. (Cat # F3165), The antibody specific to the HA tag (Cat # 923501) was purchased from Biolegend Inc. (CA, USA). The Ubiquitin antibody (P4D1, Cat # 3936S) and MG132 (Cat # 2194) were obtained from Cell Signaling Technology Inc. (MA, USA). The antibody specific for β-actin (Cat # ab8227) and biotin (Cat # ab53494) was obtained from Abcam, Inc. (UK). We purchased the antibody for CXXC5 (Cat # 16513-1-AP) from Proteintech Inc. (IL, USA). RNase A (Cat # EN0531) was obtained from ThermoFisher Scientific Inc. (Waltham, MA, USA). Goat anti-rabbit (Cat # R-05072) and goat anti-mouse (Cat # R-05071) secondary antibodies conjugated with horseradish peroxidase, and the WesternBright ECL kit (Cat # K-12045-D50) were obtained from Advansta Inc. (CA, USA). Triton X-100 (Cat # A4975) was purchased from AppliChem Inc. (Darmstadt, Germany). Protease Inhibitor, PI, (Cat # 11836170001) and Phosphatase Inhibitor, PhosSTOP, (Cat # 4906845001) were obtained from Roche Inc (Basel, Switzerland).

Pageruler Prestained Protein Ladder (ThermoFisher, Cat # 26616) or Pageruler Plus Prestained Protein Ladder (ThermoFisher, Cat # 26619) was used as the molecular mass marker.

E2 was dissolved in Ethanol (Sigma-Aldrich, Cat # 1.00983) to 10^-3^ M as the stock concentration and kept at -20°C.

### Generation of Mutant CXXC5 cDNAs

To assess the intracellular location, ubiquitination, and protein stabilities of CXXC5 mutant proteins, we used overlapping PCR^48, 92^ with primer sets (Supplementary Information, Table for Primers) specific to the 3F- or HA-CXXC5 cDNA.

### Cell growth

The growth and maintenance of MCF7 and HEK293FT cells were described previously^45, 48, 80, 104^. In brief, MCF7 cells were cultured in high glucose (4,5 g/L) containing Dulbecco’s modified Eagle’s medium (DMEM, Sartorius, Cat # 04-007-1A) supplemented with 10% fetal bovine serum (FBS, Sartorius, Cat # 01-055-1A), 1.2% L-Glutamine (Sartorius, Cat # 03-020-1B) and 1% Penicillin-Streptomycin (Sartorius, Cat # 03-031-1B). HEK293FT cells were cultured in high glucose (4,5 g/L) in DMEM supplemented with 10% FBS, 1.2% L-Glutamine, 1% Penicillin-Streptomycin, 2% Sodium Pyruvate (Sartorius, Cat # 03-042-1B) and 2% MEM Non-Essential Amino Acids Solution (Sartorius, Cat # 01-340-1B).

### Synchronization of cell cycles

To minimize the effects of mitogenic estrogens on cellular proliferation in cell cycle synchronizations, we culture MCF7 cells in DMEM supplemented with 10% charcoal dextran-stripped fetal bovine serum (CD-FBS) as described previously^45^. For this, MCF7 cells (7.5 x 10^5^) were plated in T-25 tissue culture flasks in DMEM medium supplemented with 10% CD-FBS for 72 h with media refreshing at 48 h. Cells were then incubated in the corresponding growth medium containing 10% CD-FBS without (0.01% ethanol, EtOH) or with 10^-9^ M E2 for three- to six-hour intervals up to 36 h to test the effects of E2 on cell cycle progression. At the termination, cells were collected with trypsinization. Cells were then subjected to flow cytometry, reverse transcription-quantitative polymerase chain reaction (RT-qPCR), and western blot (WB) analyses.

### Flow Cytometry

Cell cycle distribution was assessed with flow cytometry as we described previously^45, 104^. Briefly, cells were washed with 1xPBS, and pelleted. Cells were gently re-suspended in 100 µl of 2% CD-FBS containing 1xPBS, fixed, and permeabilized with ice-cold 70% ethanol overnight. After pelleting, cells were resuspended with 200 µl of 1xPBS containing 20 µg/ml propidium iodide, 200 µg/ml RNase A, and 0.4% (v/v) Triton X-100 for 30 minutes at room temperature. Cell cycle analyses were carried out with flow cytometry (NovoCyte Flow Cytometer; Agilent Technologies, CA, USA).

### RT-qPCR

Total RNA from MCF7 cells was obtained with a Macherey-Nagel™ Total RNA Isolation kit (Macherey-Nagel, Germany, Cat # 740955.50) according to the manufacturer’s instructions. In brief, cells were lysed with the kit’s lysis buffer, and the lysate was cleaned using NucleoSpin Filter columns prewashed with ethanol. The column was desalted with a desalting buffer and on-column DNase treatment was carried out for 45 minutes. The column was washed and RNA was eluted into RNase and DNase free H_2_O. Total RNA quantification and purity assessment were done with a NanoDrop 2000 (Thermo Fisher Scientific, US). Total RNA from cells was used for the cDNA synthesis using the RevertAid First Strand cDNA Synthesis Kit (Thermo-Fisher, Cat # K-1621). The SYBR Green Mastermix (BioRad, Hercules, CA, USA) and gene-specific primers (Supplementary Table S1) were used for RT-qPCR reactions on BioRad Connect Real-Time PCR, as we described previously^45, 48, 80, 104^. The relative quantitation of gene expressions was assessed with the comparative 2^-ΔΔC^_T_ method^105^. qPCR results were normalized to the geometric means^50^ of the transcripts levels of *RPLP0* (60 S acidic ribosomal protein P0) and *PUM1* (Pumilio RNA Binding Family Member 1) as reference genes in breast carcinoma cell models^51^. In RT-qPCR experiments, the MIQE Guidelines were followed^106^.

### Western Blot

Cells were lysed with RIPA lysis buffer (150 mM NaCl, 50 mM Tris-HCl pH: 8.0, 1% NP-40, 0.5% Sodium deoxycholate and 0.1% SDS) supplemented with 1xProtease Inhibitor (PI, Roche, Switzerland, Cat # 11836170001) and 1xPhosphotase Inhibitor (PhosSTOP, Roche, Switzerland, Cat # 4906845001). The lysates were then sonicated for 1 minute (100 mA amplitude) followed by a centrifugation at 16000 x g for 20 minutes. Supernatants were collected and proteins were quantified using Quick Start^TM^ Bradford Protein Assay (Bio-Rad Laboratories, USA, Cat # 5000201). Equal amounts of protein extracts (30 µg) were then subjected to SDS 10% PAGE at 100V for about 2 h. Proteins on the gel were transferred to a PVDF membrane (MilliporeSigma, USA, Cat # 3010040001) using a wet-transfer system at 100V for 70 minutes. The membrane was blocked with 5% skim milk in 0.1% Tris Buffered Saline-Tween (TBS-T), followed by the incubation with an antibody specific for Flag, HA, CXXC5, ERα, Ubiquitin, Biotin, HDAC1, or β-Actin. Membranes were incubated with an HRP-conjugated anti-mouse or anti-rabbit (Advansta Inc., USA, Cat # R-05072-500) secondary antibody. Proteins were visualized with the WesternBright ECL kit and images were captured with the ChemiDocTM Imaging System (Bio-Rad Laboratories). Quantifications of images were done using Bio-Rad Image Lab Software.

### Ubiquitin Pull-Down using the ^bio^Ub approach

HEK293FT cells were transiently co-transfected with pcDNA 3F-CXXC5 vector and pCAG expression vector bearing ^bio^Ub_6_-BirA or BirA cDNA in the growth medium supplemented with 50 µM biotin and 1 mM ATP to ensure biotinylation for 24 h. Cells were then collected and pelleted. Cells were lysed with a lysis buffer containing 3M Urea, 1%SDS, 100 mM N-Ethylmaleimide (NEM), 1xPI, and 1xPhosSTOP. Following a brief sonication, the lysate was centrifuged in 16000 x g for 20 minutes and the supernatant was collected. Protein content was quantified with Quick StartTM Bradford Protein Assay. Protein samples (3000 µg) were subjected to pull-down using NeutrAvidin Agarose beads (ThermoFisher, Cat # 29201) for 1 h at RT followed by an incubation for 2 h at 4°C. Beads were sequentially washed with wash buffer (WB) WB1 (8M Urea, 0.25% SDS in PBS), WB2 (6M Guanidine Hydrochloride in PBS), WB3 (6.4M Urea, 1M NaCl, 0.2% SDS in PBS), WB4 (4M Urea, 1M NaCl, 10% Isopropanol, 10% Ethanol, 0.2% SDS in PBS), WB5 (8M Urea, 1% SDS in PBS) and WB6 (2% SDS in PBS). Proteins bound to beads were eluted with 4X Laemmli SDS loading buffer (200 mM Tris-HCl pH 6.8, 8% SDS, 40% Glycerol, 0.8 mg/mL Bromophenol blue, and 100 mM DTT). The eluted samples were run on SDS-10%PAGE and WB was performed with the Flag, HA, CXXC5, Biotin, or Ubiquitin antibody.

### LC-MS/MS of ^bio^Ub-CXXC5 sequentially immunoprecipitated from transiently transfected HEK293FT cells

To assess the ubiquitination of 3F-CXXC5 by the use of the ^bio^Ub approach, we carried out large-scale (5 x T75 flasks) transient transfections in HEK293FT cells. Cellular extracts were subjected to immunoprecipitation with the Flag antibody followed by the biotin antibody. Precipitates were then subjected to an SDS-10% PAGE and the gel was stained with Coomassie Staining solution (0.1% Coomassie Blue G250, 1 M ammonium sulfate, 30% methanol, 3% o-phosphoric acid). Protein bands ranging between 40-100 kDA were excised. Gel samples were subjected to an in-gel digestion with trypsin at the EMBL Proteomics Core Facility (Heidelberg, Germany). Peptides were then analyzed using nanoAcquity UPLC (Waters) with a nanoAcquity trapping (nanoAcquity Symmetry C18, 5 μm, 180 μm × 20 mm) and analytical column (nanoAcquity BEH C18, 1.7 μm, 75 μm × 200 mm), which was coupled to an LTQ Orbitrap Velos Pro (ThermoFisher) using the Proxeon nanospray source. For data analysis, only peptides corresponding to **semi-trypsin digests** were considered.

### Co-Immunoprecipitation (IP)

Transiently transfected MCF7 cells for 48 h were collected and pelleted. Nuclear protein isolation was carried out with a NE-PER protein extraction kit (ThermoFisher, Cat # 78833) containing 1xPI and 1xPhosSTOP. Extracts were quantified with the Quick Start^TM^ Bradford Protein Assay. Protein lysates (500 µg) were subjected to precleaning using 25 µl Protein A (New England Biolabs, Cat # S1425S) and G (New England Biolabs, Cat # S1430S) conjugated magnetic beads for 4 hours at 4°C. After discarding beads, samples were incubated with 1 µg of the CXXC5 followed by 1 µg of rabbit IgG, or 5 µg of the Ubiquitin antibody (Cell Signaling Technology, Cat # 2729S) followed by 5 µg mouse IgG (Cell Signaling Technology, Cat # 2729) for 16 h at 4°C. Lysates were then incubated with 12 µl Protein A and G conjugated magnetic beads for 1 hour at 4°C. The beads were collected, washed with an IP wash buffer (150 mM NaCl, 10 mM HEPES pH 7.5, 10 mM MgCl2, 0.5% Igepal), and re-suspended with 2xSDS buffer (4%, w/v, SDS; 0.2%, w/v, bromophenol blue; 20%, v/v, glycerol, 150 mM DTT and 10% β-mercaptoethanol). Samples were loaded onto an SDS-10% PAGE, and WB was performed with the CXXC5 or Ubiquitin antibody.

### Assessing the stabilities of transcripts and proteins

To evaluate the stability of transcripts and proteins of CXXC5, ERα, and Cyclin D1, MCF7 cells were synchronized at G0/G1 with hormone withdrawal for 72 h. Cells were then treated without (ethanol control) or with 10^-9^ M E2 for 36 h. Cells were incubated with the growth medium containing 5µM ActD and/or 50 µg/ml CHX in the absence or presence of E2 for 0, 0.5, 1, 2, 4, and 8 h. Cellular extracts were prepared for and subjected to RT-qPCR for CXXC5, ERα, Cyclin D1 or RPLP0 transcript levels or WB using the antibody specific for CXXC5, ERα, Cyclin D1 or β-actin. To assess the degradation of CXXC5 via UPS, MCF7 cells were synchronized at G0/G1 with hormone withdrawal for 72 h. Cells were then treated without (ethanol control) or with 10^-9^ M E2 for 36 h. Cells were incubated with the growth medium containing none (DMSO control as the vehicle) or 10 µM MG132, a proteasome inhibitor, together with 50 µg/ml CHX in the absence or presence of E2 for 0, 0.5, 1, 2, 4, and 8 h. Cellular extracts were prepared for and subjected to RT-qPCR or WB using primer sets specific to CXXC5, ERα, or RPLP0 and WB with antibody specific for CXXC5, HA, ERα, or β-Actin.

### Evaluating the ubiquitination of CXXC5 and CXXC5 variants

To determine the ubiquitination of 3F-CXXC5, HA-CXXC5, and its variants, we used the ^bio^Ub approach. For this, transiently transfected HEK293FT cell extracts were subjected to NeutrAvidin Agarose beads. Bead-bound proteins were eluted with 4X SDS buffer and the eluted samples were subjected to WB using the Flag, HA, Biotin, or Ubiquitin antibody.

### Statistical analysis

Otherwise indicated all experiments were repeated for a minimum of two independent times. Results were presented as the mean ± standard deviation (SD). The significance of results with more than two biological replicates was determined using GraphPad Prism version 8.0, with a p-value of 0.05 being the limit of significance.

## SUPPLEMENTARY INFORMATION

### FIGURE LEGENDS

**Supplementary Information Fig. S1.**
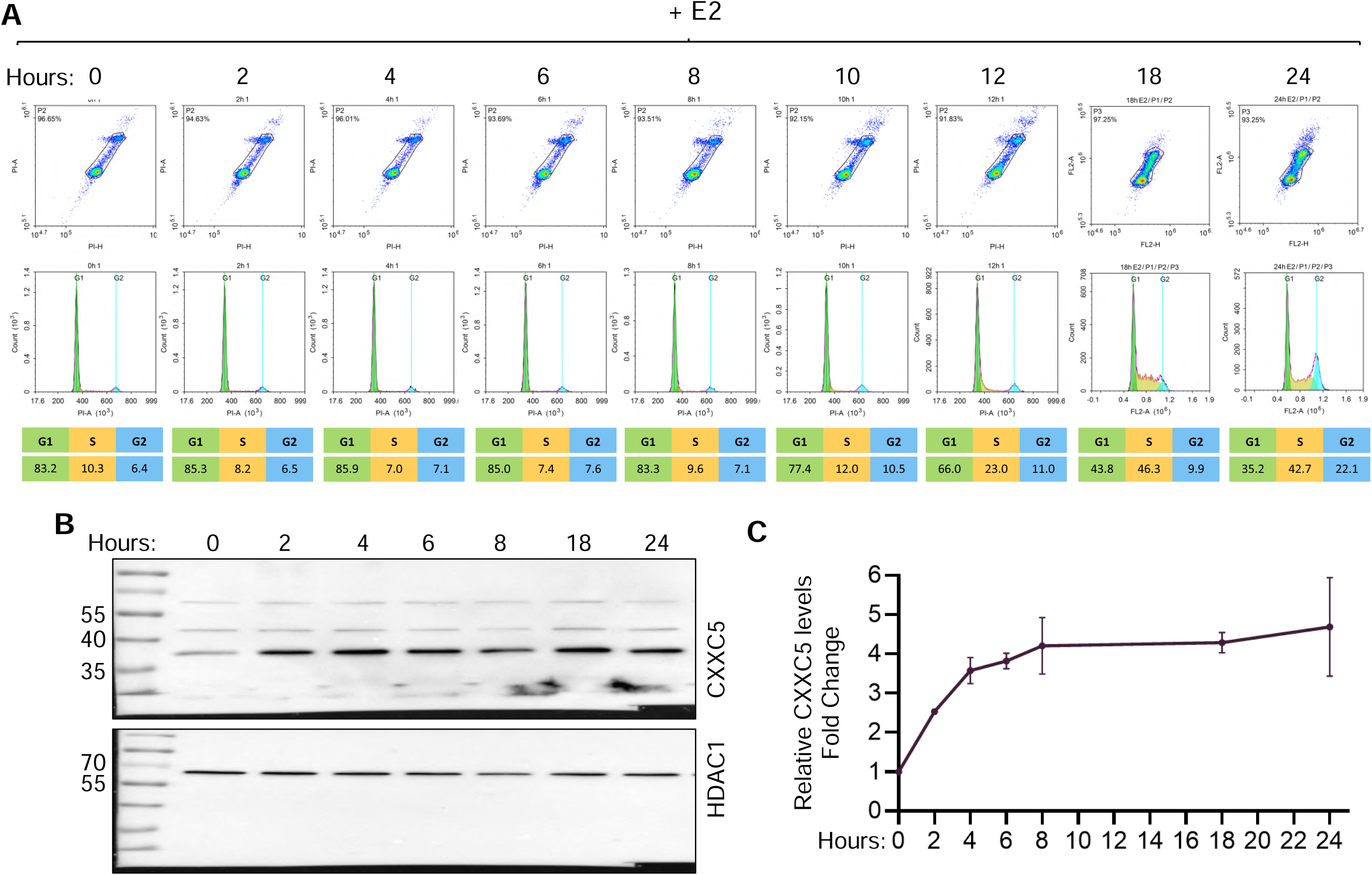
Cell cycle-dependent synthesis of CXXC5. MCF7 cells were grown in DMEM supplemented with 10% CD-FBS for 72 h with media change at 48 h. Cells were subsequently maintained in the same medium containing 0.01% ethanol (EtOH) as vehicle control or 10^-9^ M E2 for 2-6 h intervals up to 24 h. At the termination, cells were collected with trypsinization. (**A**) A fraction of cells was subjected to the flow cytometry analysis. The remaining cellular fraction was processed for and subjected to (**B**) WB analysis using the antibody specific to CXXC5, and (**C**) the results were quantified with densitometric analyses of protein levels normalized to HDAC1 which was used as the loading control. Molecular masses in kDa are indicated.

**Supplementary Information Fig. S2.**
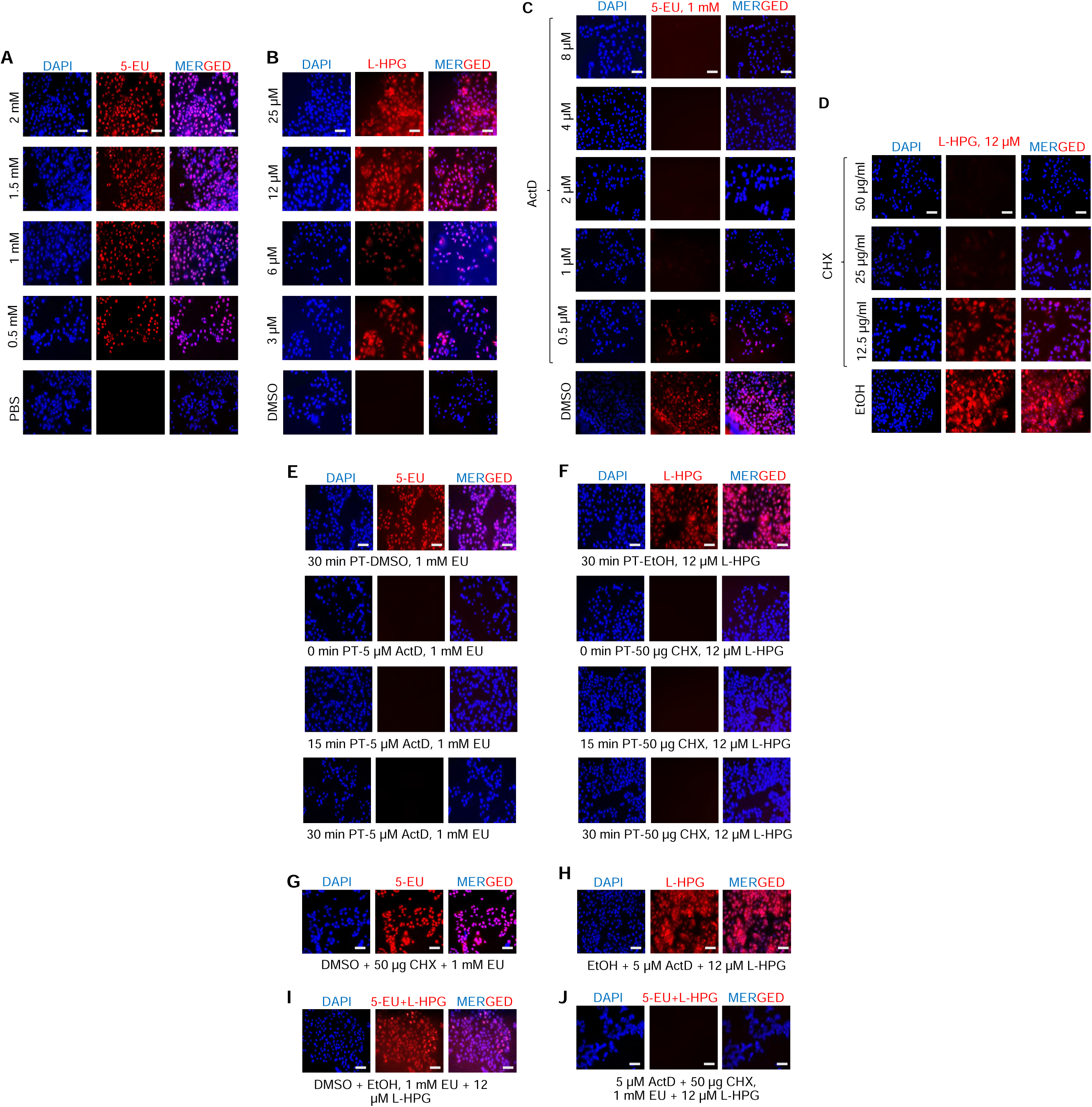
Determination of optimal concentration and treatment duration of a protein synthesis inhibitor of Cycloheximide (CHX) and RNA synthesis inhibitor of Actinomycin (ActD) in MCF7 cell. (**A**) Optimization of 5-EU concentration for the detection of nascent RNA synthesis. MCF7 cells were seeded as 15×10^3^ cells/well in 48-well plates in DMEM supplemented with 10% FBS for 48 h. The growth medium containing none (0.001% 1xPBS as the vehicle control) or 0.5, 1, 1.5, and 2 mM 5-EU for 6 h. (**B**) For the optimization of L-HPG concentration for the detection of protein synthesis, MCF7 cells (15×10^3^ cells/well in 48-well plates) grown in DMEM medium supplemented with 10% FBS for 48 h were preincubated in methionine/cysteine-free DMEM supplemented with 10 % FBS for 1 h. Cells were then incubated with the same medium containing none (DMSO, 0.001% as the vehicle control) or 3, 6, 12, and 25 μM L-HPG for 6h. (**A & B**) The cells were then subjected to a Click-IT reaction. For this, cells were washed once with 3 % BSA in 1xPBS and fixed with 3.7 % Formaldehyde in 1xPBS for 15 minutes at RT. Cells were then treated with 0.5 % Triton-X in 1xPBS for 20 minutes at RT. Cells were washed twice with 3 % BSA in 1xPBS and incubated in the Click-IT reaction buffer (20 μM Sulfo-Cy5 Azide, 100 mM sodium ascorbate, 2 mM CuSO4, and 10 mM THPTA, tris-hydroxypropyltriazolylmethylamine, in 100 mM sodium phosphate buffer, pH: 7) for 45 minutes at RT on an orbital shaker. Cells were washed twice with 3% BSA in 1xPBS, and incubated in 1xPBS containing 20 nM DAPI solution for 10 minutes at RT on an orbital shaker. After the final wash twice with 1% BSA in 1xPBS, cells were visualized in the washing solution with a fluorescent microscope. Based on the results, we concluded that 1 mM 5-EU and 12 mM L-HPG for detecting newly synthesized RNA and protein, respectively are the optimal concentrations for labeling. (**C**) To assess the optimal concentration of ActD on RNA synthesis, MCF7 cells maintained in the growth medium as described in (A) were treated without (DMSO, 0.001% as control for ActD) or with 0.5, 1, 2, 4, and 8 µM ActD in the presence of 1 mM 5-EU for 6h. Cells were then subject to the Click-IT reaction and processed for fluorescence microscopy. (**D**) For the determination of optimal CHX concentration on protein synthesis, MCF7 cells maintained and processed as described in (B) were treated without (EtOH, 0.001% as the vehicle control for CHX) or with 12.5, 25, and 50 µg/ml CHX in the presence of 12 mM L-HPG for 6 h. Cells were subject to the Click-IT reaction and processed for fluorescence microscopy. The results indicate that the optimal concentration of ActD to prevent RNA synthesis is about 5 µM; 50 µg/ml CHX is the optimal concentration to repress protein synthesis. (**E & F**) To assess whether or not the pretreatment (PT) duration of MCF7 cells with ActD or CHX is required for blocking RNA or protein synthesis, respectively, cells were preincubated without (DMSO or EtOH, as the control) or with the optimal concentration of (**E**) 5 µM ActD or (**F**) 50 µg/ml CHX for 0, 15, and 30 minutes before the addition of 1 mM 5-EU or 12 µM L-HPG. Cells were subject to the Click-IT reaction and processed for fluorescence microscopy. The results indicate that the simultaneous addition, 0 h, of ActD with 5-EU prevents RNA synthesis or of CHX with L-HPG blocks protein synthesis. These in turn suggest that the effects of ActD on RNA synthesis or of CHX on protein synthesis are immediate. (**G**) To ensure that CHX treatment does not affect RNA synthesis we co-treated cells with 1 mM 5-EU and 50 µg/ml CHX for 6h. (**H**) We also co-treated cells with 12 µM L-HPG and 5 µM ActD to examine whether or not ActD affects protein synthesis for 6h. Fluorescence microscopy results following the Click-IT reaction suggest that CHX (G) and ActD (H) at the optimal concentrations do not show cross-reactivity. (**I & J**) Co-treatment of cells with 1 mM 5-EU and 12 µM L-HPG for 6h in the presence (**J**), but not the absence (**I**), of 5 µM ActD and 50 µg/ml CHX blocks both RNA and protein synthesis. The scale bar is 50 µm.

**Supplementary Information Fig. S3.**
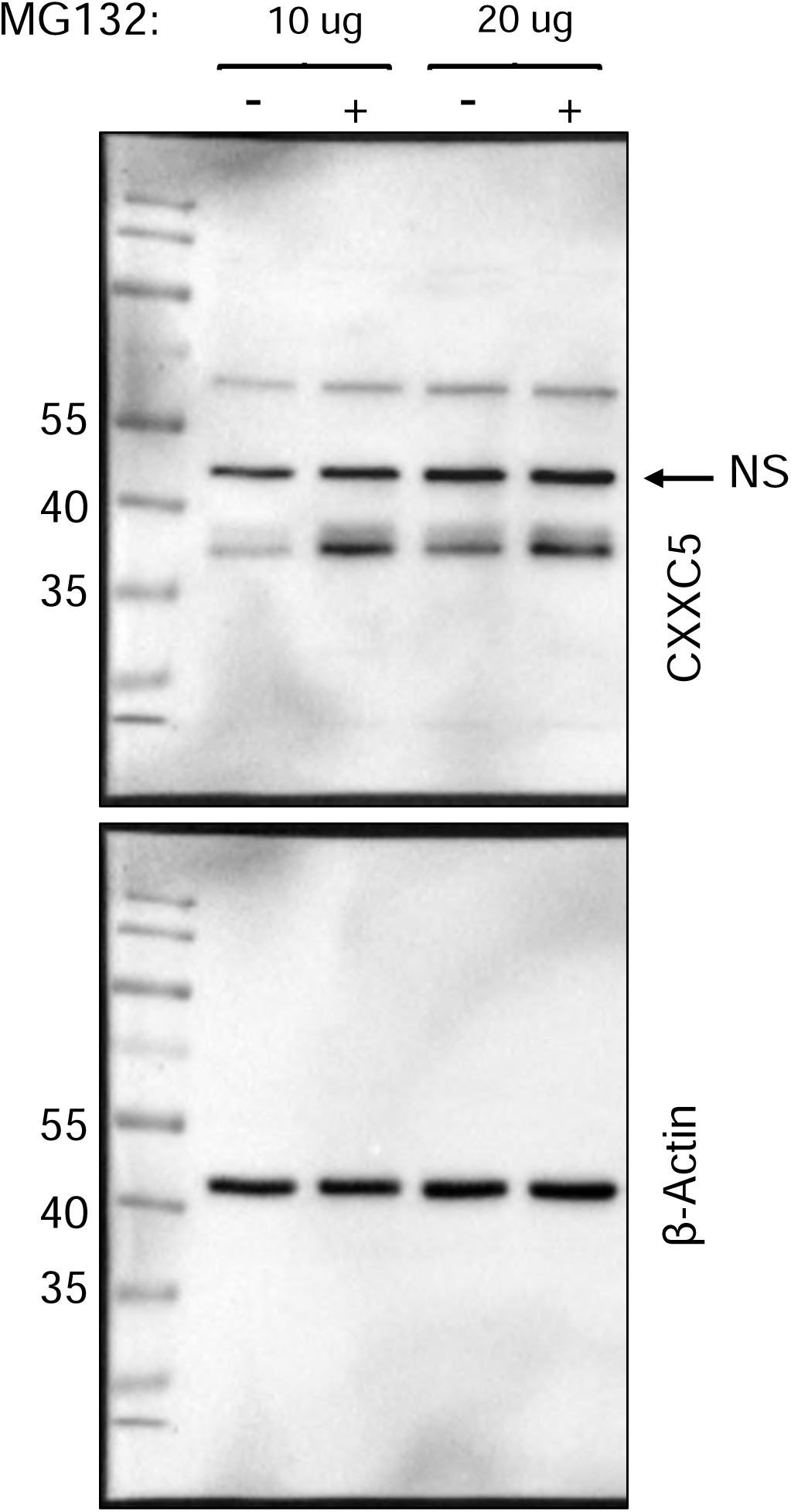
Assessing the effects of concentrations of MG132 on the stability of CXXC5. MCF7 cells grown in steady-state conditions were incubated in the growth medium without (-) or with (+) 10 or 20 µM MG132 for 4 h. Cellular extracts were subjected to WB using the CXXC5 or β-Actin antibody. Molecular masses in kDa are indicated.

**Supplementary Information Fig. S4.**
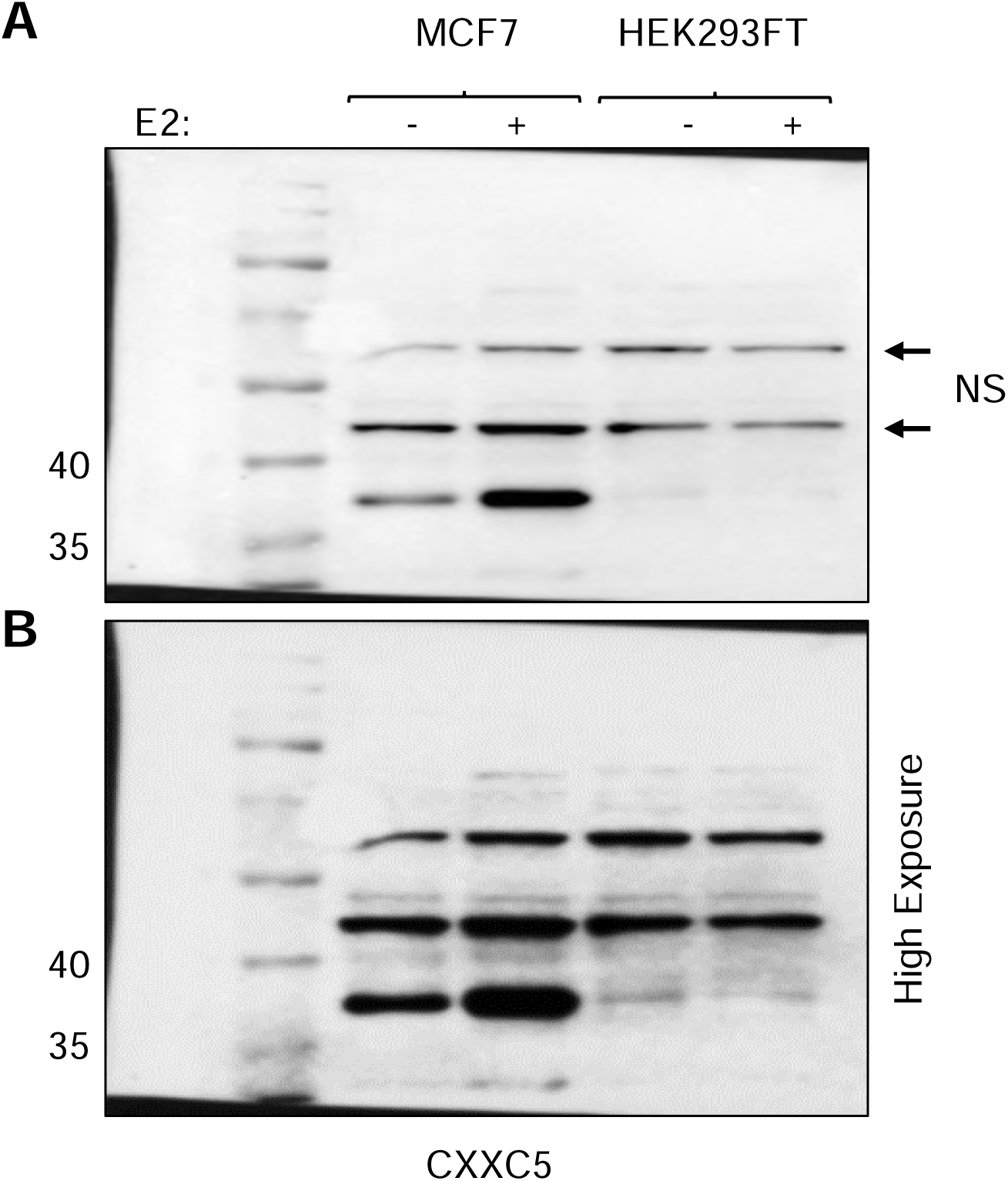
Estrogen responsiveness and CXXC5 synthesis in MCF7 and HEK293FT cells. Cells were incubated in growth media containing 10% CD-FBS for 72 h. Cells were then treated without (- E2) or with 10^-9^ M E2 (+ E2) for 24 h. Protein extracts (25 µg) were subjected to WB using the CXXC5 antibody. Molecular masses in kDa are indicated. NS indicates non-specific proteins detected by the antibody.

**Supplementary Information Fig. S5.**
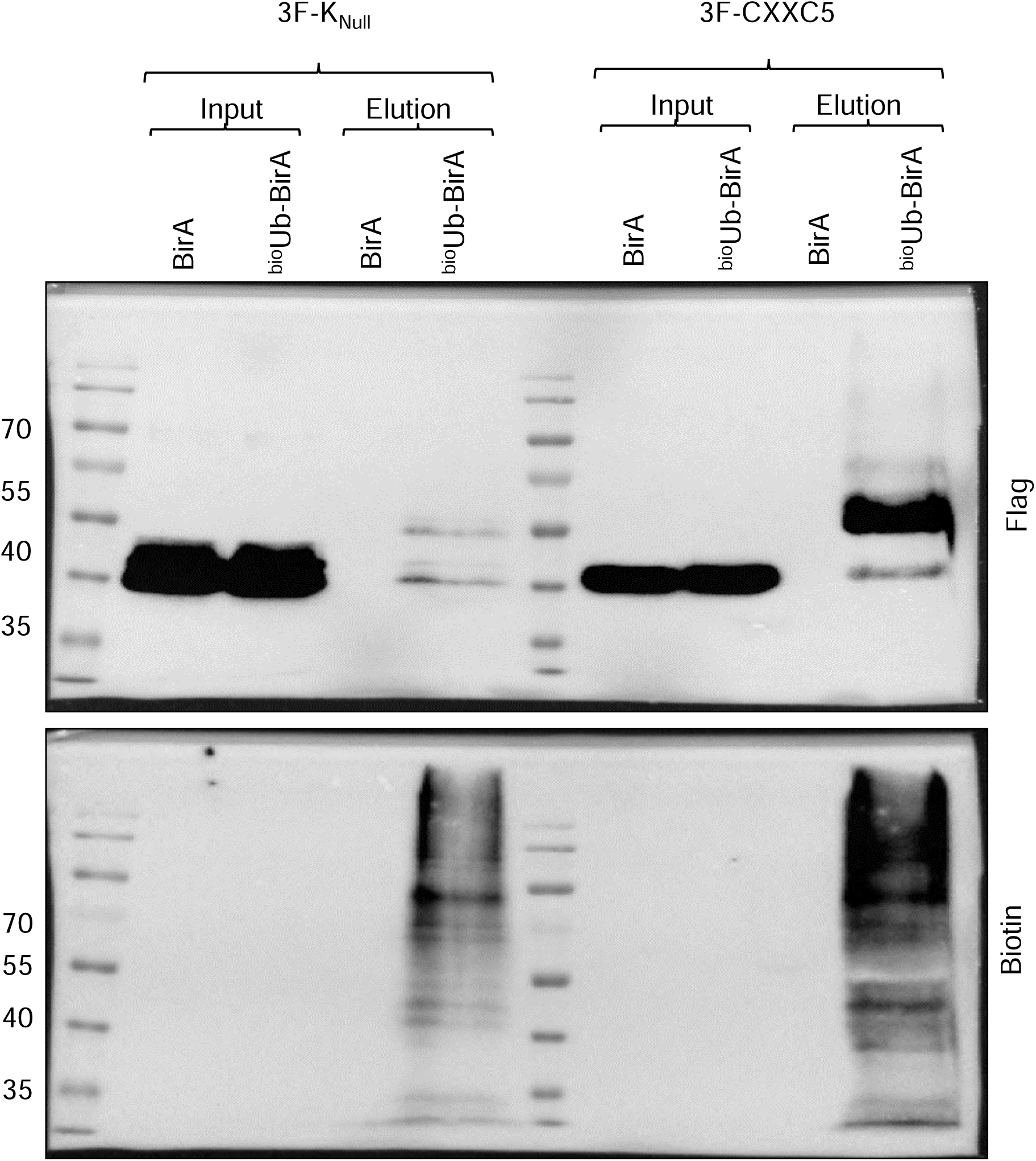
Assessing ubiquitination of K residue present in 3xFlag epitope, 3F. To assess the ubiquitination of the 3F-K_Null_ mutant in comparison with WT 3F-CXXC5 with the ^bio^Ubi approach, protein extracts of HEK293FT cells were transiently transfected with pcDNA3.1(-) expression vector bearing the 3F-K_Null_ or WT-3F-CXXC5 cDNA together with pCAG-BirA or pCAG-(^bio^Ub)_6_-BirA and incubated in a fresh growth medium containing 50 µM biotin and 1 mM ATP for 24 h. Cells were collected and extracts were subjected to NeutrAvidin-conjugated agarose beads followed by WB using the Flag or Biotin antibody.

**Supplementary Information Fig. S6.**
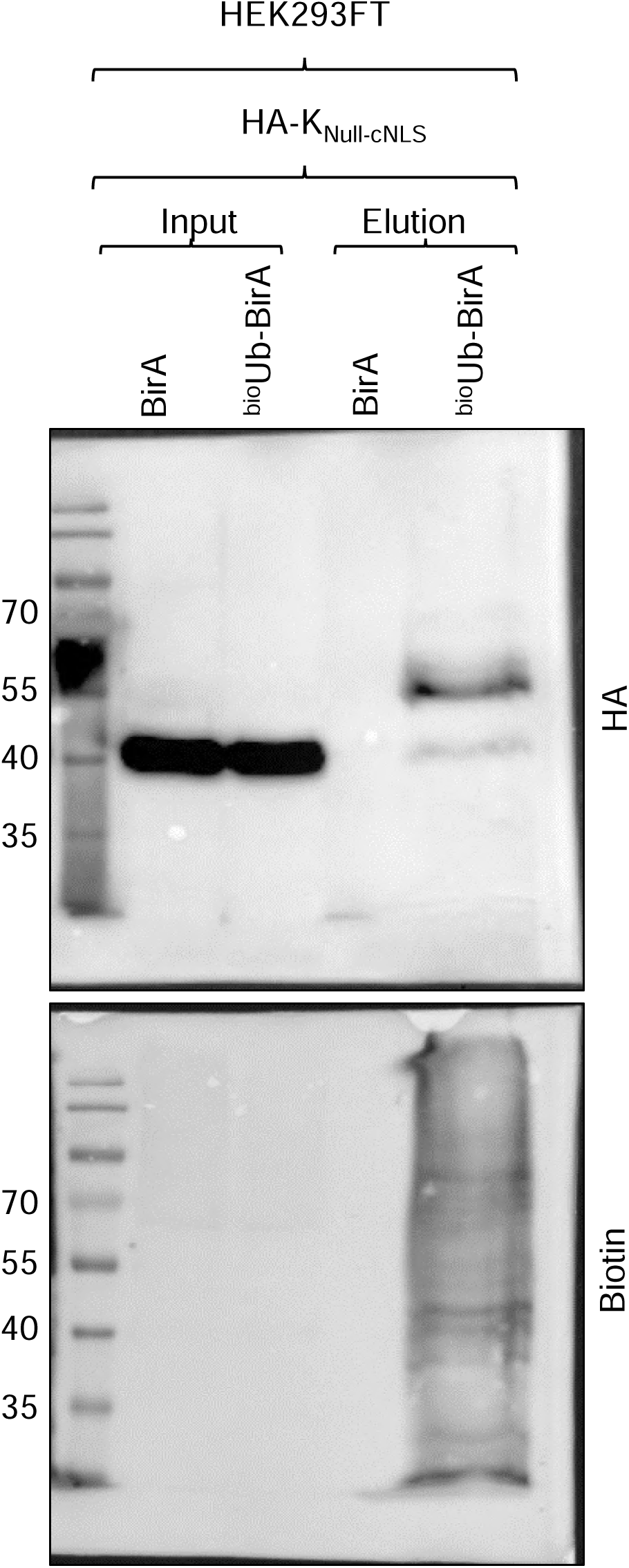
Examination of ubiquitination of K residue in the NLS of CXXC5. To assess the possible ubiquitination of the HA-K_Null_ mutant CXXC5 with the ^bio^Ubi approach, we transiently transfected HEK293FT with pcDNA3.1(-) expression vector bearing the HA-K_Null-cNLS_ cDNA together with pCAG-BirA or pCAG-(^bio^Ub)_6_-BirA expression vector and cells were then incubated in a fresh growth medium containing 50 µM biotin and 1 mM ATP for 24 h. Cellular extracts were subjected to NeutrAvidin-conjugated agarose beads followed by WB using the HA or Biotin antibody.

**Supplementary Information Fig. S7.**
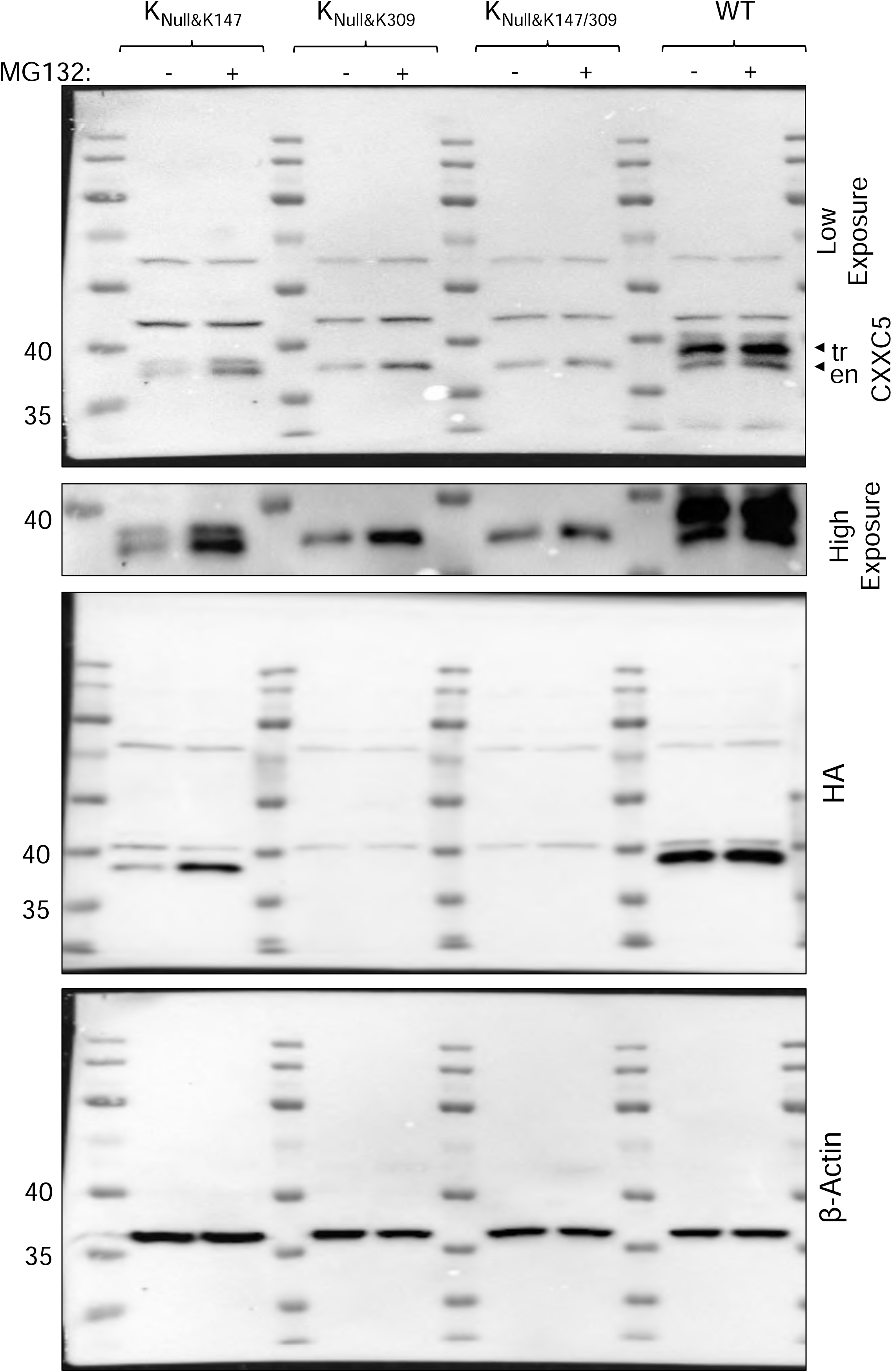
Assessing the effect of MG132 on the stability of HA-CXXC5 variants. MCF7 cells grown in steady-state conditions were transiently transfected with expression vector bearing HA-K_Null&K147_, HA-K_Null&K309_, HA-K_Null&K147/309_, or HA-WT CXXC5 cDNA for 48 h. Cells were then incubated in the growth medium without (-) or with (+) 10 µM MG132 for 4 h. Protein extracts were subjected to WB with the CXXC5 antibody and re-probed with the HA or β-Actin antibody. ‘tr’ and ‘en’ indicate transfected HA-CXXC5 and endogenous CXXC5, respectively. Molecular masses in kDa are indicated.

## ACKNOWLEDGMENTS

This study was supported by the Scientific and Technological Research Council of Turkey (TUBITAK) under grant number 121Z346 (MM). We thank TUBITAK for their support. We express our gratitude to Drs. Ugo Mayor and Juanma Ramirez at the UPV/EHU, Spain, for the kind gift of pCAG-(^bio^Ub)_6_-BirA and pCAG-BirA expression vectors. We thank Dr. Umut Şahin at the Boğaziçi University, Türkiye, for the initial guidance in ubiquitination studies. We thank Mert Kaan Çadır, Öykü Sağlık, and Sude Bildi for their technical support throughout the studies.

## AUTHOR CONTRIBUTIONS

Hazal Ayten: Conceptualization, data curation; formal analysis; investigation; methodology; writing-original draft; editing. Pelin Toker: Data curation; formal analysis; investigation; methodology; writing-original draft; editing. Gizem Turan; Investigation; methodology. Çağla Ece Olgun: Investigation; methodology. Büşra Bınarcı: Investigation; methodology. Gizem Güpür: Investigation; methodology. Pelin Yaşar: Investigation; methodology. Hesna Begüm Akman: Methodology; editing. Per Haberkant: Formal analysis; methodology; editing. Mesut Muyan: Conceptualization; supervision; funding acquisition; data curation; formal analysis; writing-original draft; editing; project administration.

## ADDITIONAL INFORMATION

### Supplemental Information

Supplementary Figures S1-S7; Supplementary Information Table for Primers, Supplementary Information Uncropped Images of Figures.

### Data Availability

This article and its supplementary information files include all data generated or analyzed during this study.

## Supplementary Information, Figures, Table and Uncropped Images

**Supplementary Information Uncropped Images Figure 1.**
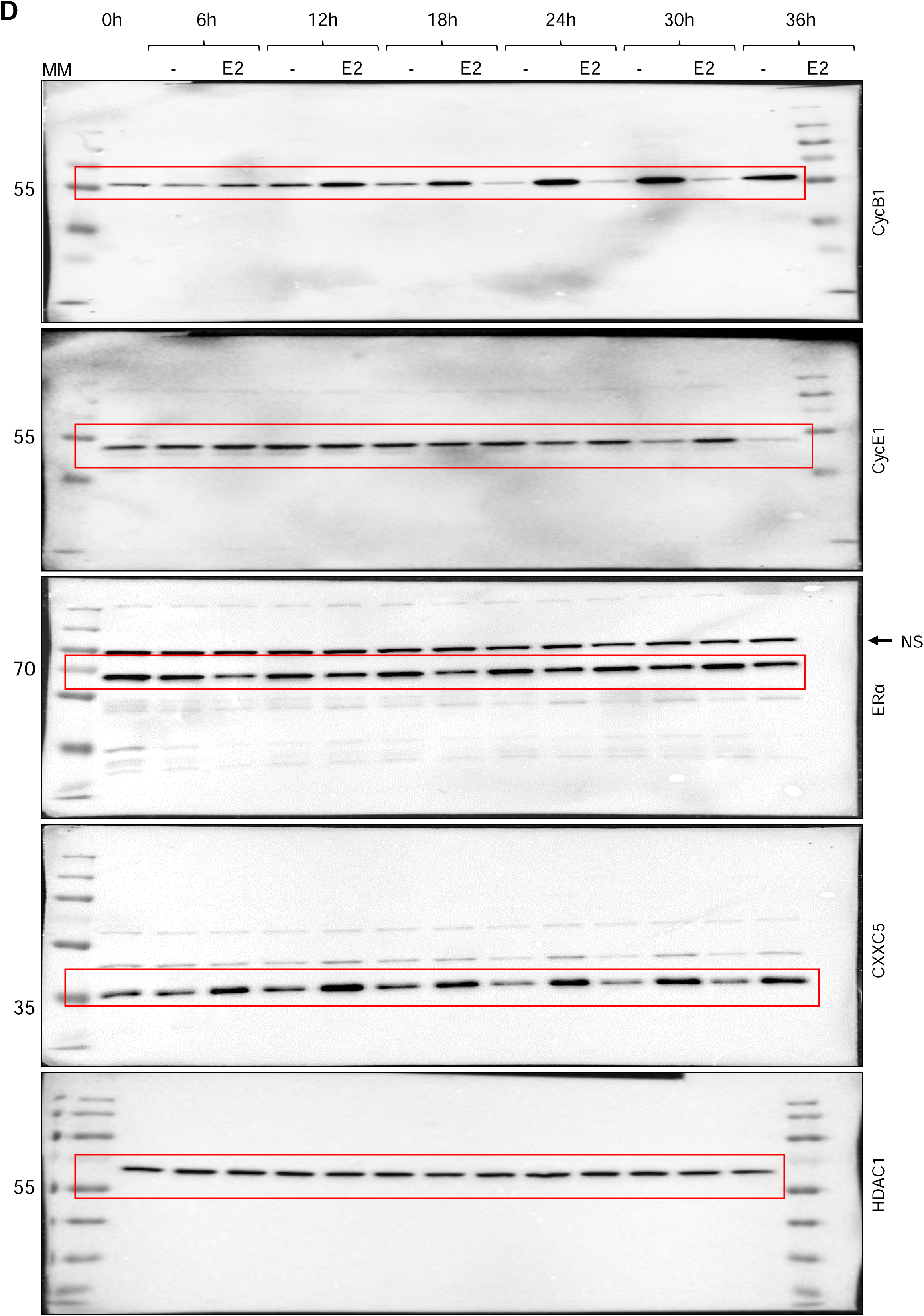

**Supplementary Information Uncropped Images Figure 2.**
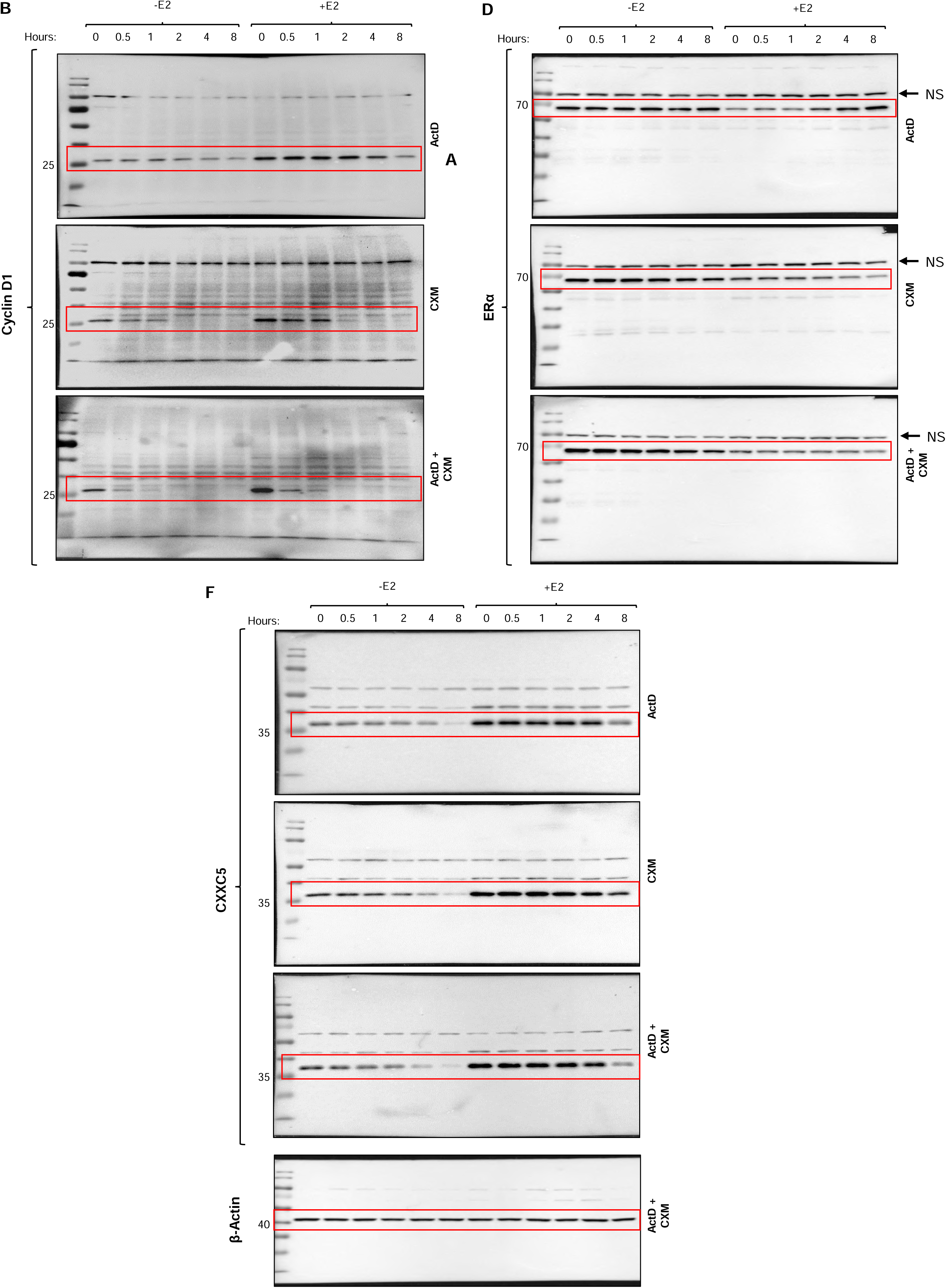

**Supplementary Information Uncropped Images Figure 3.**
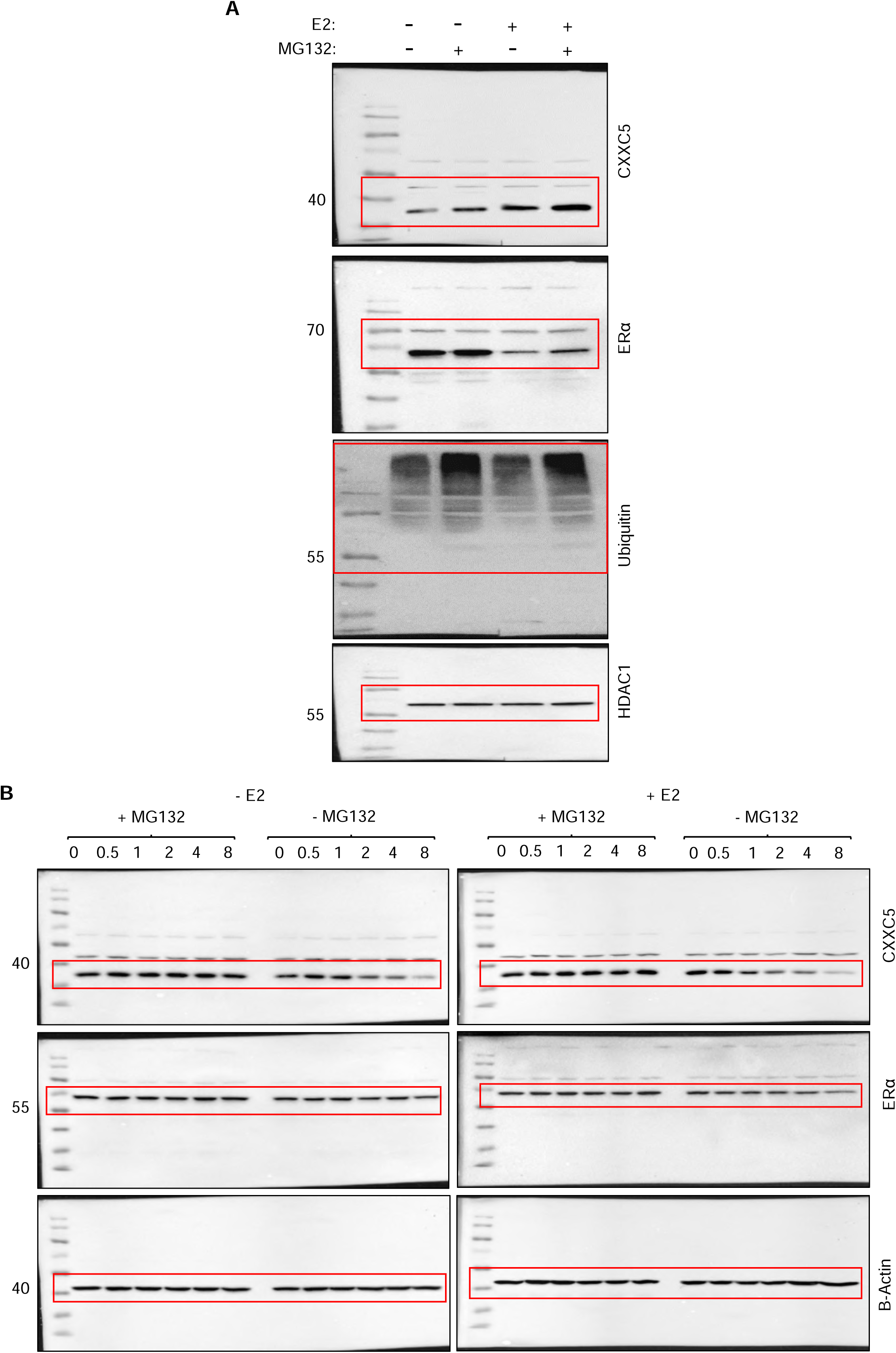

**Supplementary Information Uncropped Images Figure S4.**
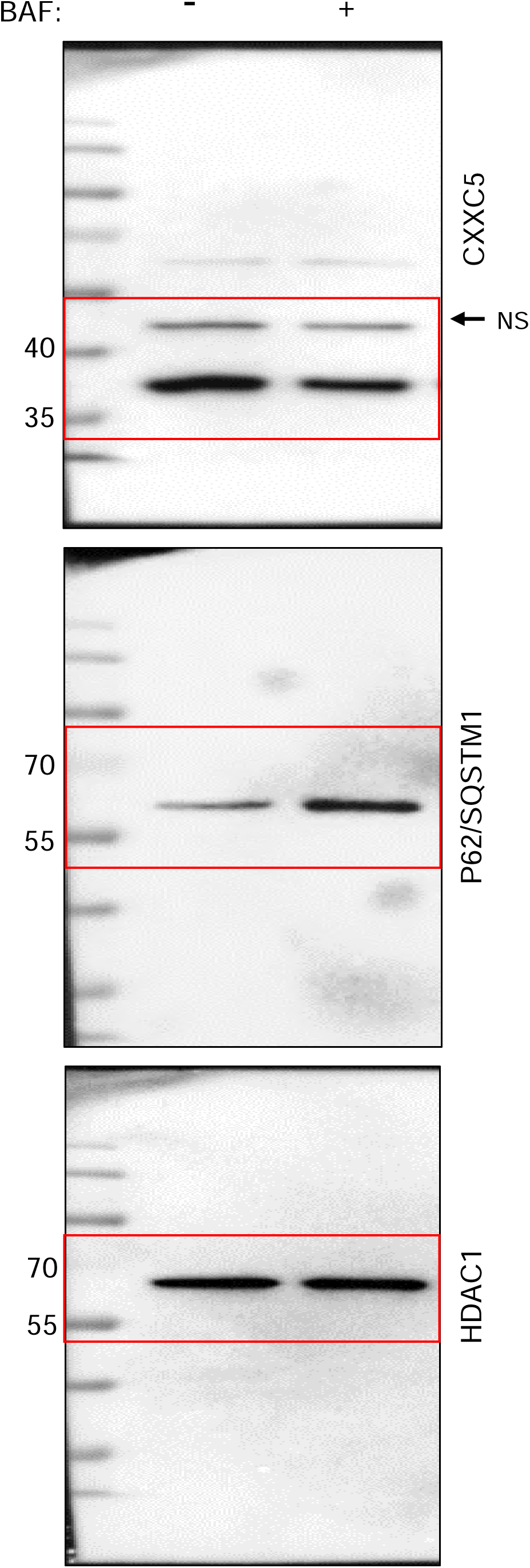

**Supplementary Information Uncropped Images, Figure 4.**
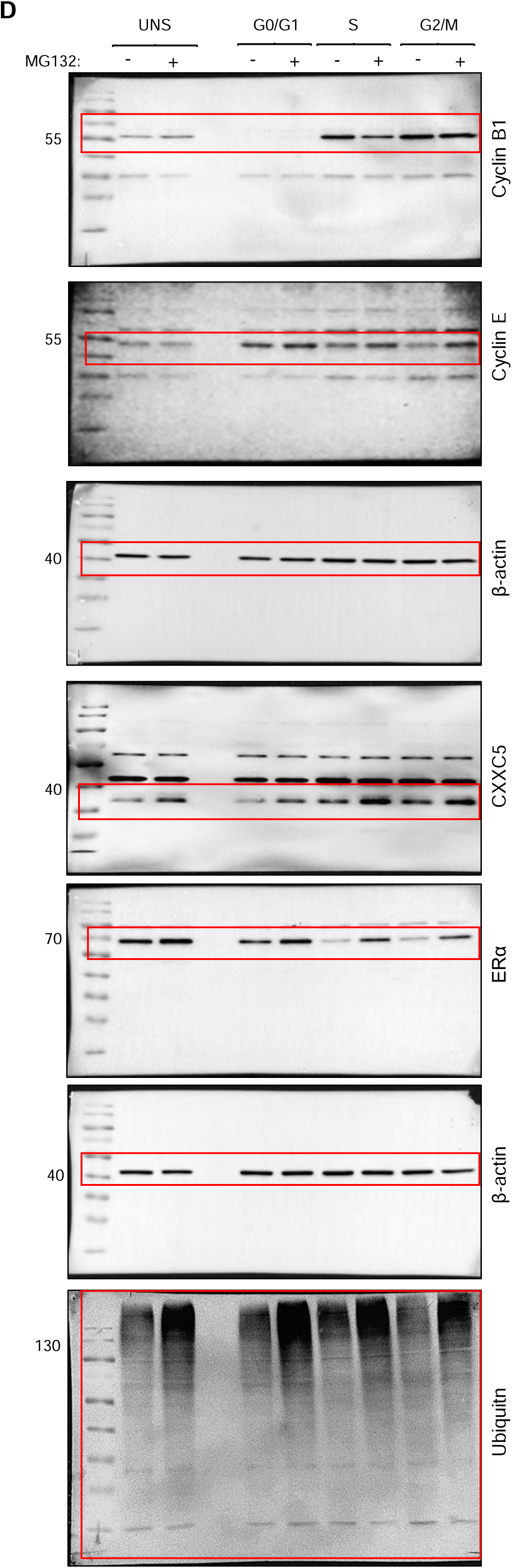

**Supplementary Information Uncropped Images, Figure 7.**
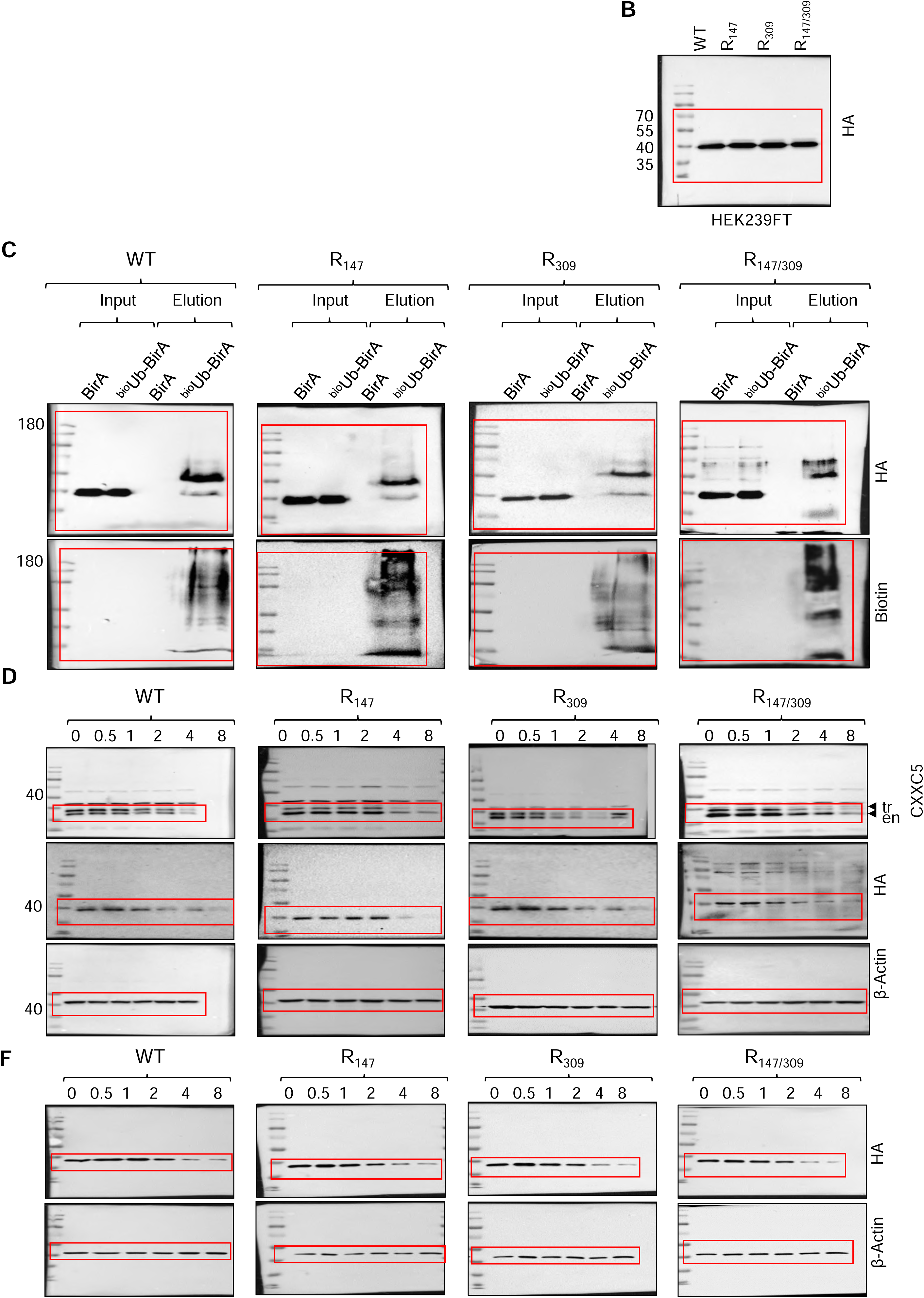

**Figure 9.**
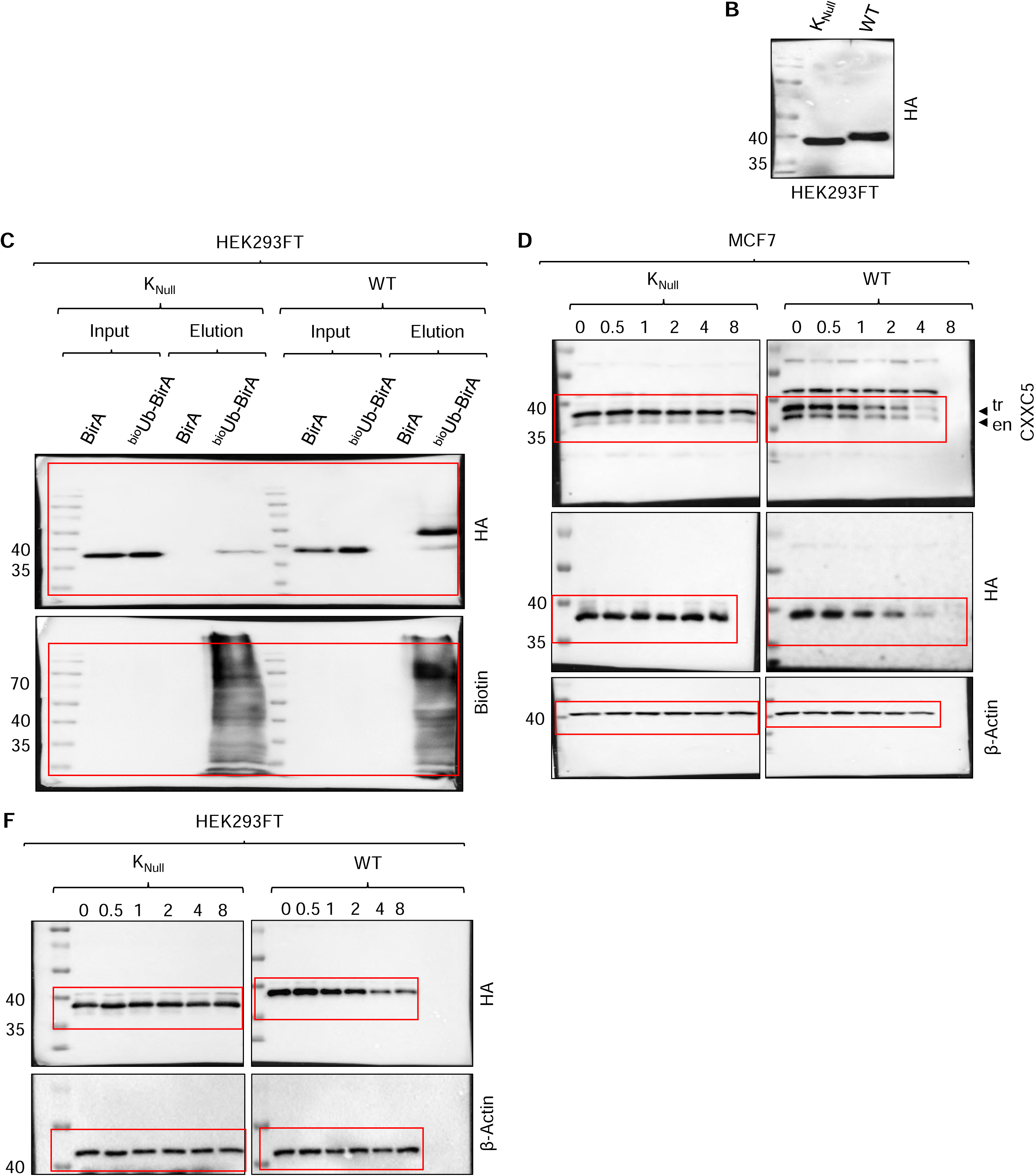

**Information Uncropped Images, Figure 10.**
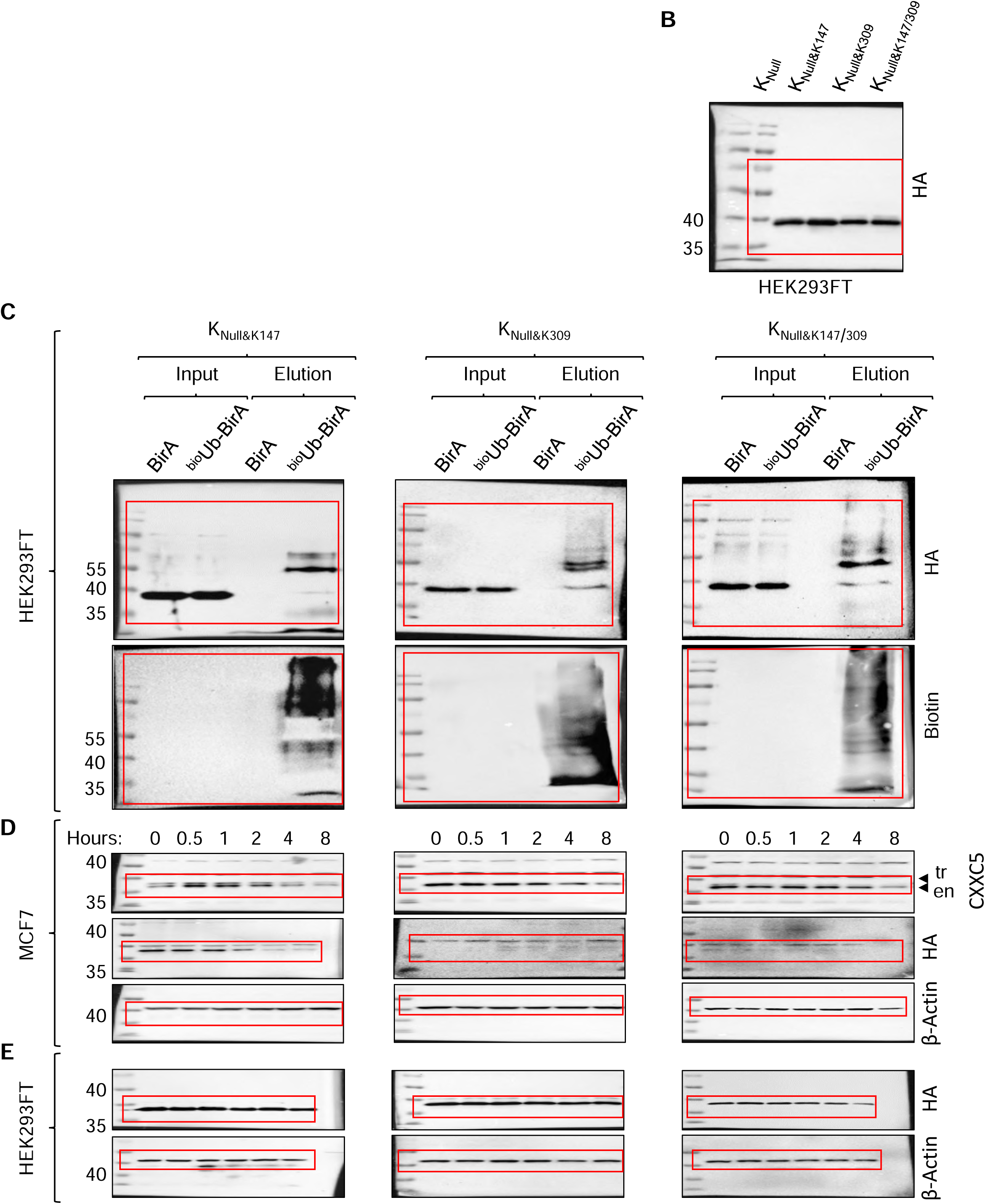

---

**Supplementary Information Table.**
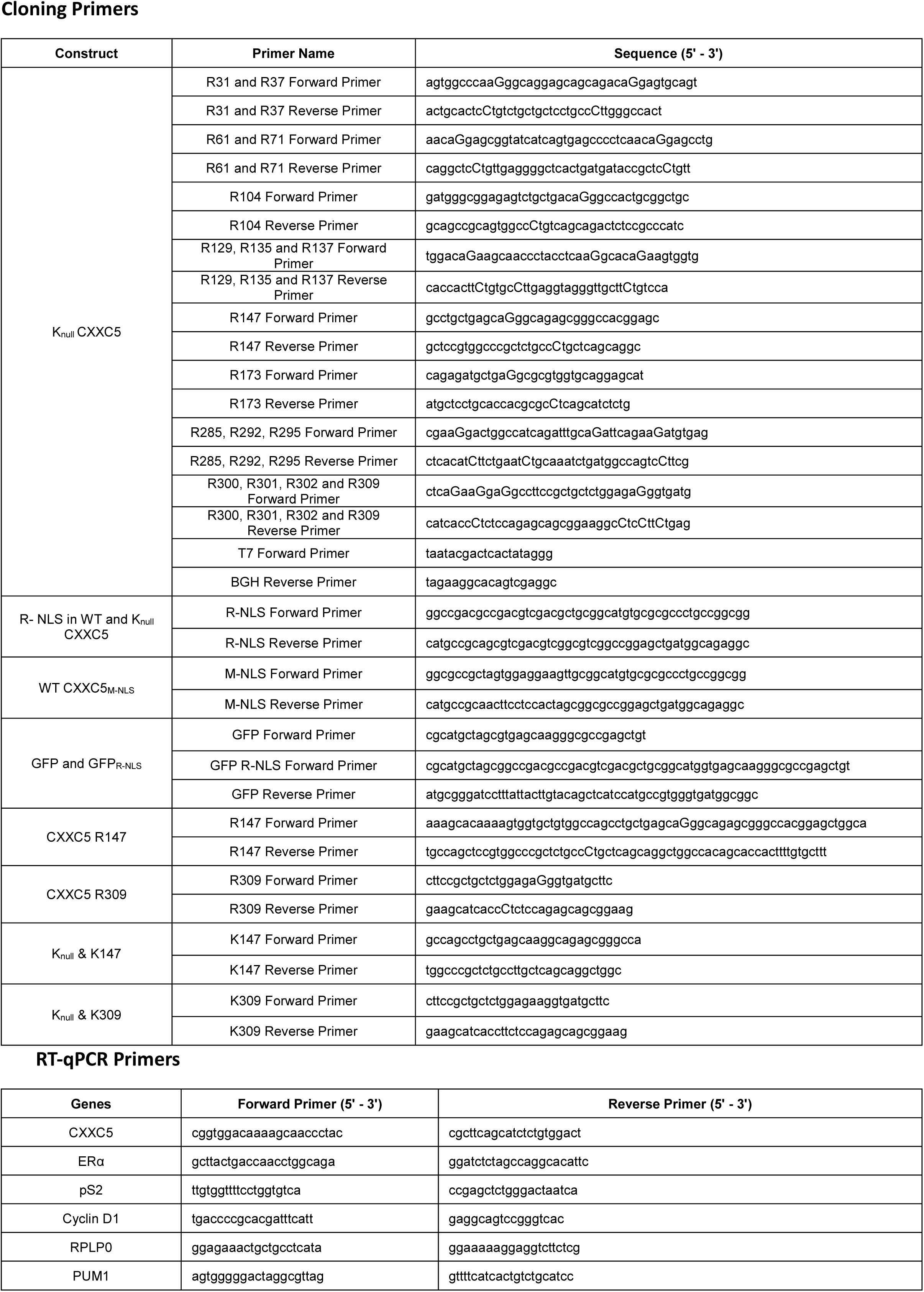
Primer Sequences Cloning Primers.

